# Topological Regulation of the Mammalian Genome by Positive DNA Supercoiling

**DOI:** 10.64898/2026.03.10.710740

**Authors:** Ashish Kumar Singh, Luis Altamirano-Pacheco, Lieng Taing, Agnès Dubois, Pablo Navarro

## Abstract

DNA supercoiling is a fundamental aspect of genome topology characterizing all DNA transactions. While negative supercoiling has been extensively characterized, positive supercoiling remains poorly understood. Using a quantitative GapR profiling system, we establish comprehensive maps of positive supercoiling across interphase and mitosis. We show that positive super-coils accumulate not only at gene ends but prominently at promoters, enhancers, insulators and loop anchors, where they are resolved by Topoisomerases. Biochemical and functional assays reveal the main sources of positive torsion: transcription generates genic supercoils, while R-loops and Cohesin drive accumulation at regulatory elements, topologically associating domains and their boundaries. During mitotic chromosomal compaction, Condensins establish a global wave of positive supercoiling that largely homogenizes the genome, yet promoters with rapid post-mitotic reactivation retain elevated torsion and R-loops. These findings establish positive DNA supercoiling as a form of topological memory that links DNA mechanics to transcriptional control, genome architecture and epigenetic inheritance.

## Introduction

Genomic DNA undergoes substantial torsional stress throughout the cell cycle and across all domains of life (Vino-grad et al. 1965; Duprey and Groisman 2021). The torsional stress manifests in the formation of DNA supercoils by changing the DNA twist and intertwined loops called plectonemes (Gilbert and Marenduzzo 2025). DNA supercoiling has been primarily associated with the strand separation activity of DNA and RNA polymerases during transcription and replication (Wu et al. 1988; Yu and Dröge 2014). As polymerases translocate along the DNA, they induce over-twist in the front and undertwist in their wake, a phenomenon described in the twin-domain supercoil model as positive and negative supercoiling, respectively (Liu and Wang 1987; Gilbert and Marenduzzo 2025).

DNA supercoiling regulates several fundamental biological processes. DNA melting by negative supercoils is known to promote transcription initiation, binding of gene regulatory factors, and the formation of non-B DNA structures (Parvin and Sharp 1993; Stolz et al. 2019). Positive supercoiling, on the contrary, inhibits transcription elongation and destabilizes DNA-bound factors and nucleosomes (Gartenberg and Wang 1992; Sheinin et al. 2013; Teves and Henikoff 2014). Excessive local DNA supercoiling can propagate to neighboring regions, causing inadvertent activation or repression of adjacent regulatory regions and genes. For instance, positive supercoiling may lead to eviction of transcription factors or DNA melting by negative supercoils can lead to R-loops, RNA–DNA hybrids with a displaced single-stranded DNA and typically enriched at active promoters and enhancers (Massé and Drolet 1999; Chedin and Benham 2020; Patel et al. 2023; Figueroa-Bossi et al. 2024; Morao et al. 2025). Thus, dedicated mechanisms such as DNA Topoisomerases I (Top1) and II (Top2) control and resolve DNA supercoiling in concert with RNA Polymerase II (Pol II) at active genes but also at boundaries of topologically associating domains (TADs) generated by Cohesin-mediated loop extrusion (Vos et al. 2011; Kouzine et al. 2013; Naughton et al. 2013; Baranello et al. 2016; Uusküla-Reimand et al. 2016). In line with their key role in regulating DNA topology, Topoisomerases are essential in transcriptional regulation, DNA loop formation during interphase and genome condensation during mitosis (Neguembor et al. 2021; Hirano 2025).

All three Structural Maintenance of Chromosomes (SMC) complexes – Cohesin, Condensin, and Smc5/6 – have been shown to alter DNA topology during loop extrusion in vitro on naked DNA (Kim et al. 2022; Jeppsson et al. 2024; Davidson et al. 2025). Notably, Condensins were shown to induce positive supercoiling in the presence of Top2. This activity is predicted to help compact mitotic chromosomes by inducing positive supercoils (Kimura and Hirano 1997; Baxter and Aragón 2012). However, even if yeast plasmids exhibit positive supercoiling during anaphase upon depletion of Top2 (Baxter et al. 2011), a direct demonstration of genome-wide positive supercoiling in mitosis is still lacking. Positive supercoiling may also contribute to chromatin compaction during interphase and promote enhancer-promoter contacts (Benedetti, Dorier, and Stasiak 2014; Fu et al. 2024). Introduction of positive supercoiling during simulations recapitulates many features of the interphase 3D-genome, suggesting that TADs may be positively supercoiled (Benedetti, Dorier, Burnier, et al. 2014; Racko et al. 2019). Nevertheless, whether TADs and other DNA loops are indeed positive supercoiled remains to be established in vivo.

Current methods for mapping DNA supercoiling primarily rely on DNA intercalants, have low resolution and are hardly quantitative (Kouzine et al. 2013; Naughton et al. 2013; Teves and Henikoff 2014; Hall et al. 2025; Yao et al. 2025). Recently, the bacterial protein GapR was repurposed to detect positive supercoils in bacteria, yeast, and higher eukaryotes (Guo et al. 2021; Diman et al. 2024; Longo et al. 2024). Despite this critical advance, our understanding of the mechanisms driving positive DNA supercoiling during the cell cycle remains completely lacking. Here, we establish an inducible, spike-in normalized GapR system in mouse Embryonic Stem (ES) cells and show that positive supercoils are pervasively generated by R-loops and Cohesins at regulatory elements, loop anchors and TADs. During mitosis, we show that Condensins trigger a wave of positive super-coiling genome-wide, erasing most differences observed in interphase. However, promoters displaying fast reactivation dynamics during mitosis maintain particularly high levels of positive supercoiling, associated with R-loops persistence. We propose that positive supercoiling plays determinant roles in gene regulatory processes and their mitotic inheritance.

## Results

### Genome-wide map of positive supercoiling in mouse embryonic stem cells

To map positive supercoiling, we established doxycycline (dox)-inducible ES cell lines expressing similar levels of FLAG-tagged wild-type GapR and mutant (GapRMUT), which can bind DNA but does not accumulate at positive supercoils (Guo et al. 2021) (Figure S1A, B). Dox induction of GapR for 16 h did not alter growth, ES cell self-renewal or gene expression (Figure S1C-E). GapR localized exclusively to the nucleus, whereas GapRMUT showed diffused signal throughout the cell (Figure S1F). To determine genome-wide GapR localization, we performed quantitative ChIP-seq using a spike-in approach with E. coli expressing GapR. We observed GapR but not GapRMUT signal throughout the genome, accumulating over highly transcribed genes such as histone clusters (Figure 1A, top), and under the form of focal peaks reminiscent of epigenomic marks at regulatory elements (Figure 1A, bottom). To explore this systematically, we performed Pol II ChIP-seq and ranked all genes accordingly (Figure 1B, left). Monitoring GapR signal across these ordered genes clearly showed a prominent and focal enrichment of GapR at the TSS (Transcription Start Site) across all transcribed genes and a broader signal at the TES (Transcription End Site) of highly transcribed genes (Figure 1B, right). Similar patterns were also detected after ranking genes by their RNA levels (Figure S1G). While the Pol II-correlated enrichment at TES was expected and consistent with the twin domain supercoil model, the specific enrichment at active promoters was not, given that these regions are supposedly negatively supercoiled. Even if this observation recapitulates an earlier study (Longo et al. 2024), we aimed at exploring this feature in more detail.

**Fig. 1.**
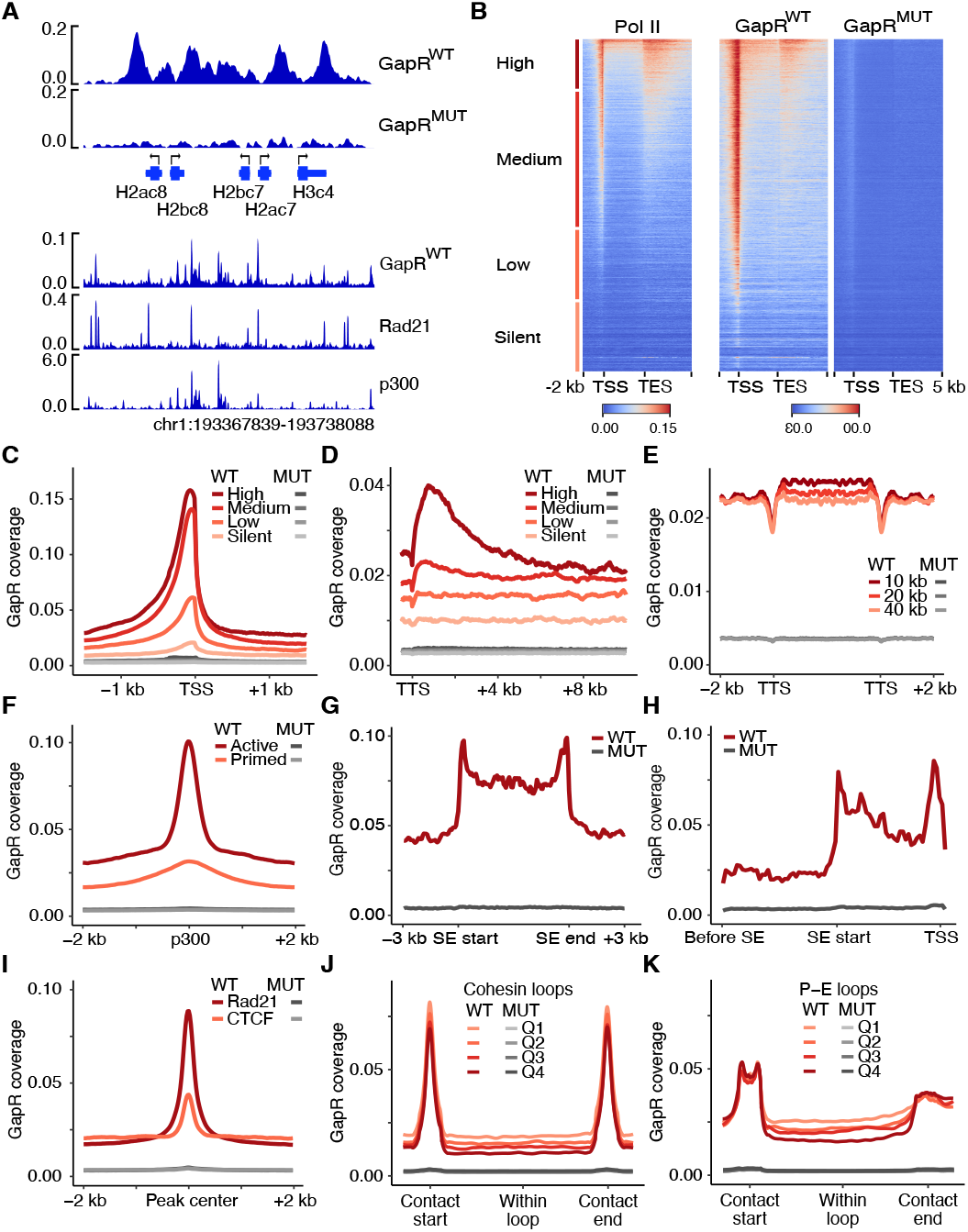
Widespread positive supercoiling at regulatory elements. (A) Illustrative coverage of E. coli spike-in normalized GapRWT/MUT-FLAG ChIP signal at the histone locus (top) and at a representative region showing GapRWT enrichment overlapping Rad21 and p300 binding sites outside genic regions (bottom, mm10 coordinates). (B) Average heatmaps of Pol II (n=2, N20 antibody), GapRWT and GapRMUT signal (n=3) across all coding genes, sorted by global Pol II occupancy per gene. Color scales represent Pol II and spike-in normlaized GapR ChIP signal. (C) Average profiles of GapRWT/MUT ChIP signal centered on TSS and stratified by the four Pol II categories shown in B. (D) Same as C, but for 10 kb long regions starting at TES. (E) GapRWT/MUT signal between expressed convergent genes grouped by the distance between two convergent TES. (F) Average profiles of GapRWT/MUT ChIP centered on p300 summits for active and primed enhancers (Cruz-Molina et al. 2017). (G) Average binned profiles of GapRWT/MUT at SEs. (H) Average binned profiles of GapRWT/MUT ChIP signal centered on the SE start and spanning an equidistant region with its target gene and the upstream region (“Before SE”). (I) Average profiles of GapRWT/MUT ChIP signal centered on regions binding Rad21 and CTCF (labelled Rad21) or CTCF only (labelled CTCF). (J) Average binned profiles of GapRWT/MUT within Cohesin DNA loops. Signal was aligned to start-end of interacting loop anchors and divided into quartiles based on the loop size: Median Q1 = 84 kb, Q2 = 144 kb, Q3 = 190 kb and Q4 = 244 kb. (K) Same as J but for promoter-enhancer (P-E) loops. Median P-E loop sizes for each quartile were Q1 = 55 kb, Q2 = 119 kb, Q3 = 215 kb and Q4 = 251 kb.

Firstly, we divided genes into high, medium, low and silent based on Pol II levels (Figure 1B, left) and assessed GapR binding around their TSS and TES. GapR binding scaled with Pol II and was sharply localized immediately upstream of the TSS (Figure 1C). In contrast, GapR binding downstream of TES extended up to 6 kb for high Pol II genes and 2 kb for medium Pol II genes (Figure 1D). Positive supercoiling also enriched between pairs of expressed, convergent genes and scaled with the distance between genes (Figure 1E, S1H). Secondly, we focused on GapR peaks and found 78k with a median peak size of 600 bp, while GapRMUT yielded no detectable peaks, further confirming the specificity of our experimental system. In accord with the gene-centered analysis, GapR peaks were enriched near promoters relative to their genome distribution (Figure S1I). ChromHMM-based annotation (Pintacuda et al. 2017) confirmed GapR localization to active promoters, but also at enhancers, bivalent chromatin and insulator regions, while being depleted in intergenic and heterochromatin regions (Figure S1J). These correlations were confirmed by establishing the average GapR binding profiles across ChromHMM categories (Figure S1K). We conclude, therefore, that GapR accumulates at expected regions of positive supercoiling (TES) but also at regions supposedly characterized by negative supercoiling such as promoters, enhancers and insulators.

### Regulatory elements and their loops are positively super-coiled by distinct mechanisms

Given the enrichment of GapR peaks over regulatory elements, which are often characterized by accessible chromatin, we asked whether GapR binding is merely biased to open chromatin, even though GapRMUT is not recruited at these sites. Comparison of GapR ChIP-seq with ATAC-seq revealed that while virtually all ATAC peaks are enriched for GapR, it binds substantially more regions (56k) than ATAC peaks (Figure S1L), demonstrating that GapR does not simply recognize open chromatin. After excluding the possibility of a mere bias to accessible regions we focused on enhancers. Consistent with Pol II activity at enhancers (Heinz et al. 2015; Cruz-Molina et al. 2017), we observed strong GapR binding at active enhancers, including super-enhancers (SEs), whereas primed enhancers with low Pol II activity showed minimal GapR signal (Figure 1F, G). Notably, GapR signal near SEs exhibited a pronounced directional pattern, oriented toward the target gene contacted by the SE, while the DNA upstream of the SEs lacked such pattern (Figure 1H). This suggests that directional positive supercoiling of DNA between enhancers and promoters may foster their 3D contacts. Accordingly, we examined GapR binding near TAD boundaries co-occupied by the loop extruding factor Cohesin and the insulator protein CTCF (Merkenschlager and Nora 2016; Liu et al. 2021). Regions binding Rad21 and Smc1, core subunits of Cohesin, and the insulator protein CTCF blocking loop extrusion, showed a strong signal whereas regions bound only by CTCF had weaker GapR signal (Figure 1I, S1M), suggesting that Cohesin may generate positively supercoiling at TAD boundaries. Moreover, within loops categorized as either TADs or promoter-enhancer (P-E) and promoter-promoter (P-P) interactions (Hsieh et al. 2022) displayed GapR enrichment, especially within small loops and scaled with loop size (Figure 1J, K, S1N). Thus, positive supercoiling appears to spread within loops but dissipates over longer distances.

Finally, to test whether transcription and Topoisomerases modulate positive supercoiling, we used chemical inhibitors targeting Top1 (camptothecin), Top2 (etoposide), transcription initiation (triptolide) or elongation (flavopiridol; Figure S2A), with no measurable effects on cell viability (Figure S2B). Indeed, Topoisomerases inhibition led to increased GapR binding over the TSS and TES of Pol II bound genes compared to silent genes that showed no change (Figure 2A top, B, C). Consistent with increased GapR signal at TES, we also observed elevated GapR binding between convergent genes following Top1 and Top2 inhibition (Figure S2C). Similarly, active enhancers (Figure 2D), SEs (Figure 2E, F), and TAD boundaries (Figure 2G), exhibited increased GapR binding upon Topoisomerases inhibition, not only within but also between 3D anchoring regions (Figure 2F, H, I, S2D). Thus, Topoisomerases effectively resolve positive supercoiling throughout the genome. In contrast, transcription inhibition led to decreased GapR binding only at TES and between convergent genes, with the strongest effects seen at highly transcribed genes and after inhibition of elongation (Figure 2A, bottom), but had no effect at TSS (Figure 2J, K, L) or at any other regulatory elements and their connecting DNA loops (Figure S2E-K).

**Fig. 2.**
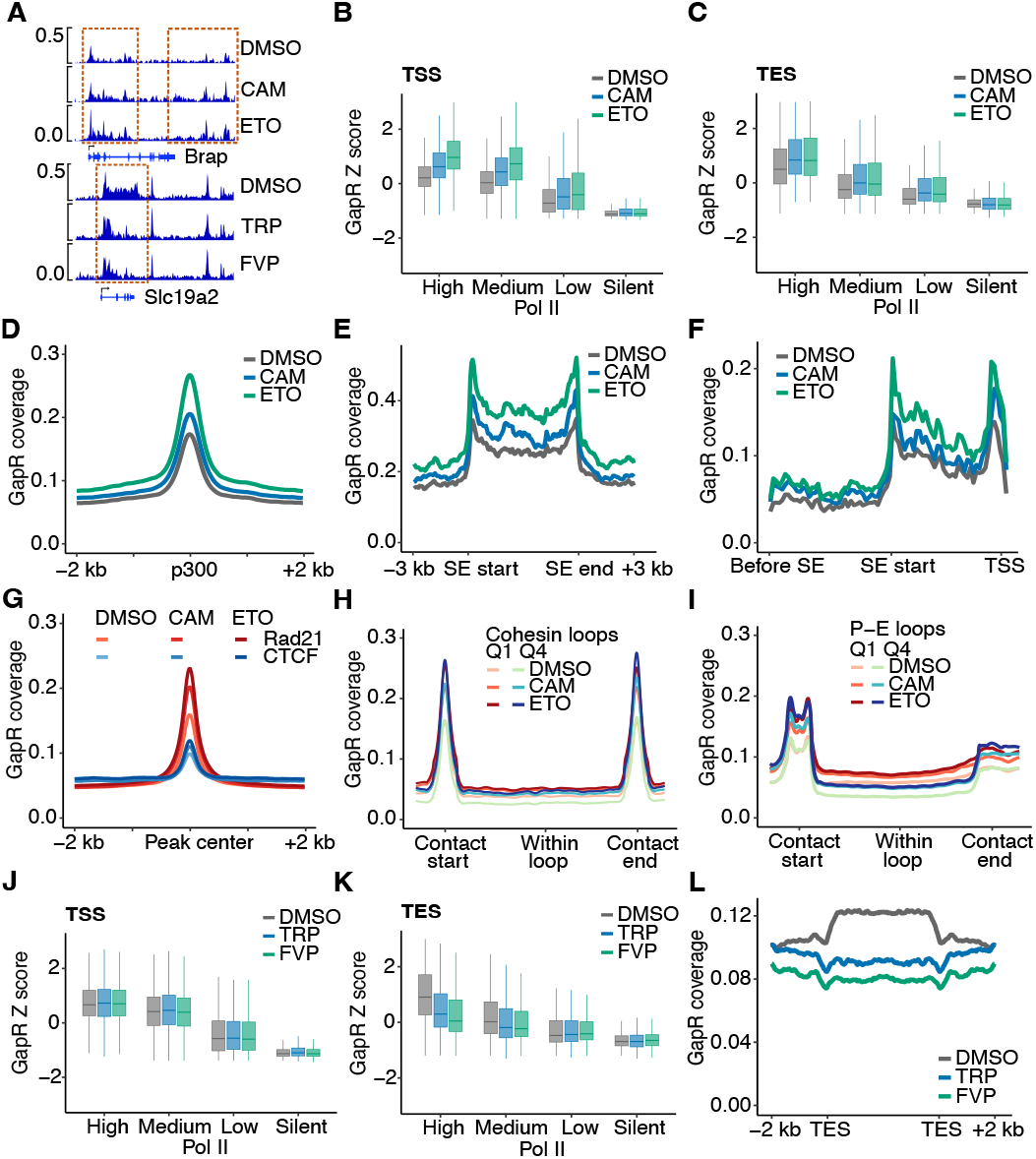
Topoisomerases resolve positive supercoiling genome-wide, while transcription generates positively supercoiling primarily near TES. (A) Illustrative coverage of spike-in normalized GapR signal of a representative gene after treatment with DMSO, topoisomerases or transcription inhibitors. Regions responding to inhibition are highlighted. (B) Boxplots quantifying GapR signal (n=2) from TSS across the four Pol II categories shown in Figure 1B using the [-1, 0.2 kb] interval. (C) Same as B but for signal downstream of TES (5kb). (D) Average binned profiles of GapR signal near active enhancers centered on p300 summits. (E) Average binned profiles of GapR signal at SE aligned to SE start and SE end coordinates. (F) Average binned profiles of GapR centered on the SE start and spanning an equidistant distant with its target gene and the upstream region. (G) Same as D but for centered on regions binding Rad21 and CTCF (labelled Rad21) or CTCF only (labelled CTCF). (H, I) Average binned profiles of GapR within the smallest (Q1) and largest (Q4) cohesin (H) and P-E (I) loops following topoisomerase inhibition. (J) Same as B, but for cells treated with transcription initiation (TRP) and elongation (FVP) inhibitors. (K) Same as C, but for cells treated with transcription inhibitors. (L) Average binned profiles of GapR signal between expressed convergent genes grouped by the distance between two convergent TES of 10 kb for cells treated with transcription inhibitors.

These results suggest that transcription generates positive supercoiling primarily at TES, contrasting with the transcription-dependent generation of negative supercoiling at TSS (Yao et al. 2025). The accumulation of positive supercoiling at regulatory elements may thus depend on other molecular sources, which we elucidate below.

### R-loops generate localized positive supercoil at regulatory sites

Because transcription inhibition did not alter GapR binding near key regulatory sites, we asked whether alternative mechanisms may generate positive supercoiling at these genomic loci. We reasoned that at these sites positive supercoiling might be generated as a compensatory torsion to the generation of negative supercoiling. One common feature of active regulatory elements are R-loops (Li et al. 2016), which induce DNA melting and may trigger adjacent compensatory positive supercoiling. As shown before in yeast, positive supercoiling accumulates near R-loops (Guo et al. 2021).

Using high-resolution mapping of R-loops in ES cells (Wulfridge and Sarma 2021), we observed that R-loops accumulate immediately upstream of the GapR peak at TSS and, strikingly, that their abundance correlates with both Pol II occupancy and GapR signal (Figure 3A, B). To validate this observation, we performed R-loop CUTTag (Kaya-Okur et al. 2019) and ranked all R-loop peaks by decreasing signal. We observed a proportional scaling of GapR at these sites (Figure 3C), supporting the idea that R-loops may contribute to the generation of local positive supercoiling.

**Fig. 3.**
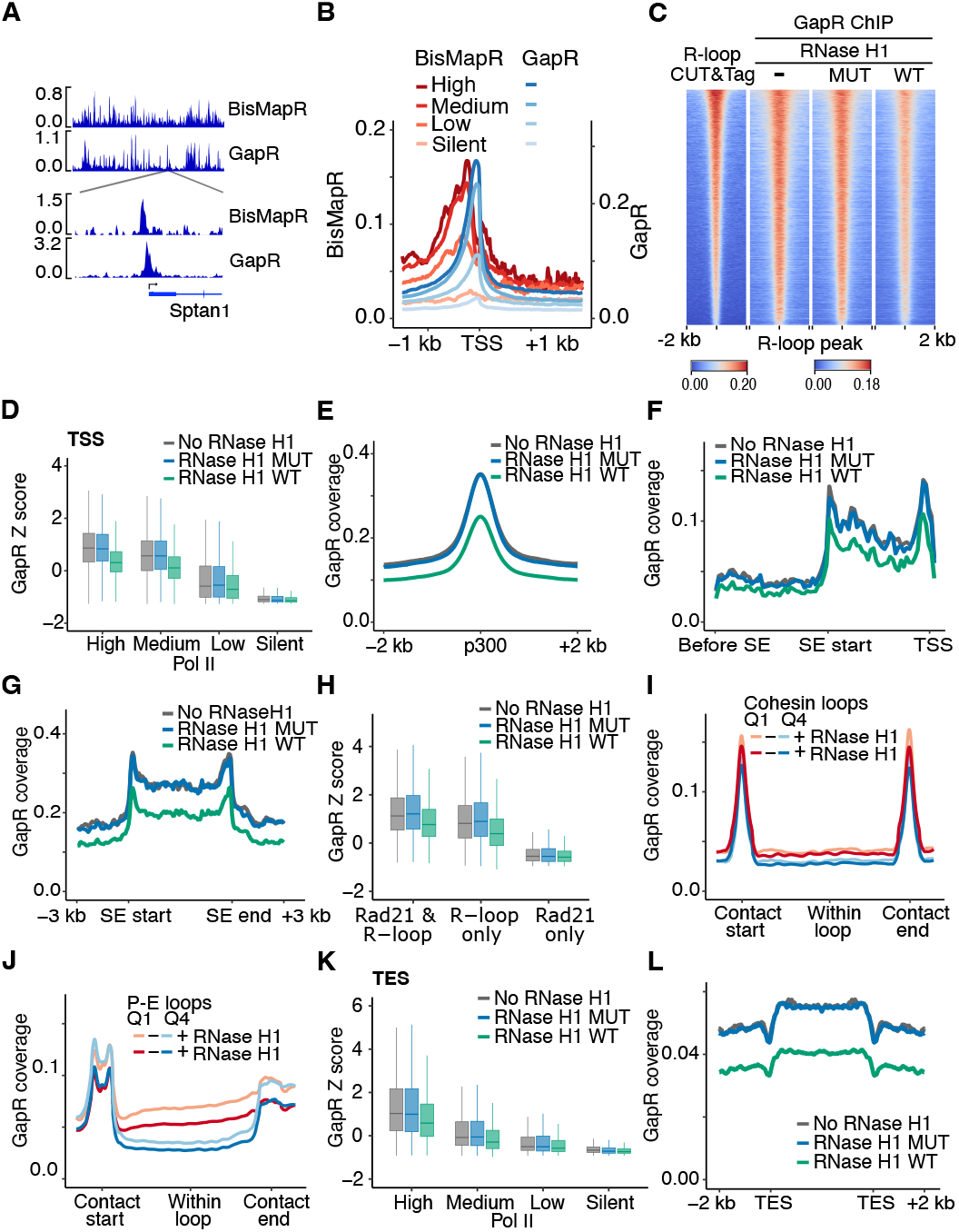
R-loops drive localized positive supercoiling at gene regulatory elements. Illustrative coverage of BisMapR and GapR signal at a representative locus (chr2:29581064-30197931) and a zoomed in view of a gene showing BisMapR and GapR signal near TSS. (B) Average profile of BisMapR R-loop and GapR signals centered on TSSs and stratified by the four Pol II categories shown before in Figure 1B. (C) Average heatmaps of peaks identified with S9.6 antibody-based R-loop CUTTag, showing signal for R-loops (n=3) and GapR (n=2) in ES cells expressing no RNase H1, MUT RNase H1 or WT RNase H1, sorted by R-loop intensity per peak. (D) Boxplots quantifying GapR signal (n=2) from TSS across the four Pol II categories using the [-1,0.2 kb] interval in cells lacking RNase H1 or expressing WT or MUT RNase H1. (E) Average profiles of GapR signal at active enhancers centered on p300 summits in cells lacking RNase H1 or expressing WT or MUT RNase H1. (F) Average binned profiles of GapR signal at SE aligned to SE start and SE end coordinates. (G) Average binned profiles of GapR centered on the SE start and spanning an equidistant distant with its target gene and the upstream region. (H) Boxplots quantifying GapR signal at peaks co-occupied by Rad21 and R-loops, compared with peaks containing only R-loops or Rad21 and CTCF, in cells lacking RNase H1 or expressing WT or MUT RNase H1. (I, J) Average binned profiles of GapR within the smallest (Q1) and largest (Q4) cohesin (I) and P-E (J) loops following RNase H1 expression. (K) Same as D, but for GapR signal from TES to TES+5000 bp. (L) Average binned profiles of GapR between convergent genes (10 kb distance) in cells lacking RNase H1 or expressing WT or MUT RNase H1.

To deplete R-loops, we generated cell lines expressing WT RNase H1 or catalytic-dead RNase H1MUT (Crossley et al. 2021) in our ES lines expressing GapR (Figure S3A). Expression of RNase H1 did not change GapR expression (Figure S3B) but led to a significant reduction in binding throughout loop peaks, whereas RNase H1MUT did not affect GapR signal (Figure 3C), suggesting a general role of R-loops in generating positive supercoils. Consistent with our hypothesis, RNase H1 markedly reduced GapR binding across all classes of regions that we had found insensitive to Pol II inhibition, such as TSS (Figure 3D), enhancers (Figure 3E), SEs and the regions linking them with their targets (Figure 3F, G), as well as TAD boundaries (Figure 3H) and over all classes of loops (Figure 3I, J, S3C). However, the dependence on R-loops was not found identical within categories. At TSS, we observed that only High and Medium Pol II promoters displayed a strong reduction (Figure 3D). At TAD boundaries, we could observe that only the minority of boundaries overlapping with R-loops peaks (Figure S3D; Wulfridge et al. 2023; Zhang et al. 2023), were sensitive to RNaseH1 depletion (Figure 3H). In line with this, GapR signal at general TAD loops was much less responsive to RNaseH1 activity than at loops associated with promoters and enhancers (Figure 3I, J, S3C). Moreover, and in agreement with the presence of R-loops near TES of highly expressed genes (Promonet et al. 2020), we also observed reduced GapR binding at the 3’ end of highly transcribed genes and between pairs of expressed convergent genes (Figure 3K, L).

Together, these results demonstrate that R-loops represent a novel mechanism contributing to the pervasive presence of positive supercoiling throughout regulatory elements, especially at promoters and enhancers and, to a lower extent, at TAD boundaries.

### Cohesin induces positive supercoiling at TAD boundaries and within loops

Neither transcription inhibition nor R-loop depletion altered positive supercoiling at the majority of TAD boundaries, suggesting that another factor may be responsible. We hypothesized that the Cohesin complex, enriched at these sites, may induce positive supercoiling through its DNA extrusion activity, as suggested by the directional positive supercoiling observed between SEs and their targets. In agreement with this, we observed that accumulation of Rad21 throughout the genome is quantitatively correlated with GapR binding (Figure 4A). To directly test the generation of positive supercoiling by Cohesin, we endogenously fused Rad21 to the degradation tag (dTAG) sequence (Nabet et al. 2018) (Figure S4A). Tagging Rad21 with dTAG reduced its expression before induction of its degradation (Figure S4B) but did not affect the growth (Figure S4C), self-renewing capacity (Figure S4D) or GapR expression (Figure S4E), suggesting that the hypomorphic nature of Rad21-dTAG is compatible with normal functions. Induction of Rad21 depletion was, moreover, fast and acute upon induction (Figure S4B) and, after 4h, did not significantly affect the cell cycle (Figure S4F).

**Fig. 4.**
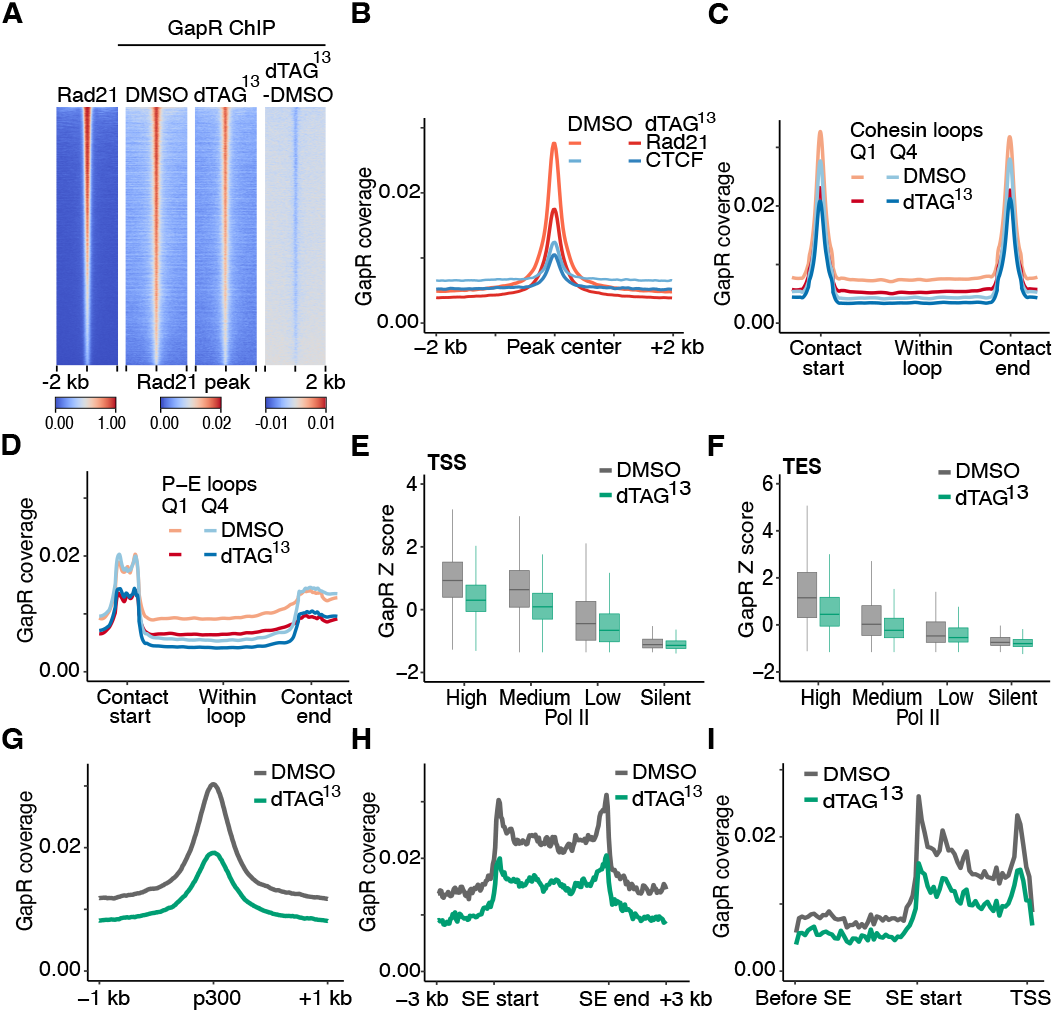
Cohesin promotes positively supercoiling at TAD boundaries and in DNA loops. (A) Average heatmaps of Rad21 peaks showing Rad21 (left; (Liu et al. 2021)) and GapR (n=2) ChIP signal in Rad21-dTAG ES cells, treated with DMSO or dTAG13 for 4 h, sorted by Rad21 intensity. On the right, the difference in signal between dTAG and DMSO treatment is shown. (B) Average profiles of GapR ChIP signal centered on regions binding Rad21 and CTCF (labelled Rad21) or CTCF only (labelled CTCF). (C) Average binned profiles of GapR signal within the smallest (Q1) and largest (Q4) cohesin loops in Rad21-dTAG cells treated with DMSO or dTAG13 for 4 h. (D) Same as C for P-E loops. (E) Boxplots quantifying GapR signal (n=2) from TSS across the four Pol II categories shown in Figure 1B using the [-1, 0.2 kb] interval. (F) Same as E but for signal downstream of TES (5kb). (G) Average binned profiles of GapR signal near active enhancers centered on p300 summits. (H) Average binned profiles of GapR signal at SE aligned to SE start and SE end coordinates. (I) Average binned profiles of GapR centered on the SE start and spanning an equidistant distant with its target gene and the upstream region.

Depletion of Rad21 led to a reduction in GapR binding throughout all Rad21 sites (Figure 4A), suggesting that Cohesin does indeed induce positive supercoiling at TAD boundaries. In contrast, CTCF-only sites lacking Rad21 did not show a significant difference (Figure 4B), underscoring the specificity of the observed effect. Next, we tested if Cohesin also generates positive supercoiling in DNA loops. At TADs, Rad21 depletion reduced GapR binding across all loop sizes, with the most pronounced decrease observed in the smallest Q1 loops (Figure 4C, D, S4G), strongly contrasting with the minor effects observed upon transcription inhibition or R-loops degradation. Moreover, and in accord with the binding of Cohesin at active TSS, TES, enhancers and SEs (Hnisz et al. 2013; Busslinger et al. 2017), Rad21 depletion also led to a reduction of GapR binding across these elements. At TSS and TES, the reduction was particularly prominent for genes with high Pol II levels (Figure 4E, F, S4H). At enhancers (Figure 4G) and SEs (Figure 4H), the reduction in GapR binding was strong and was associated with the reduction of signal throughout the regions looping with their targets (Figure 4I). Interestingly, Rad21 depletion resulted in a greater decrease in GapR signal in the direction of SE-to-promoter contact compared to the upstream region (Figure 4I, S4I), suggesting that Cohesin-driven positive supercoiling may directly facilitate enhancer-promoter communications.

Overall, these findings suggest that cohesin contributes to the formation of positive supercoiling at its binding sites, including TAD boundaries. Cohesin may further induce positive supercoiling within DNA loops, with the signal dissipating over distance, supporting a model where supercoiling spreads over large distances and contributes to the establishment of the 3D genome.

### Condensins trigger a global wave of positive supercoiling during mitosis

After establishing the prevalent presence of positive supercoiling throughout the genome and identifying the mechanisms underlying its generation, we reasoned that GapR could also be used to address a 25 years-old hypothesis: the prevalent positive supercoiling of DNA during mitosis (Kimura and Hirano 1997). To unravel the positive supercoil status of mitotic DNA, and after confirming normal GapR expression in prometaphase-arrested ES cells (Figure S5A, B), we observed that GapR, but not GapRMUT homogeneously coated mitotic chromosomes, indicating a strong enrichment for positive supercoiling (Figure 5A). Using spike-in normalized GapR ChIP-seq, we observed an increase in GapR binding across the entire mitotic genome relative to the levels measured in asynchronous cells (Figure 5B), with a substantial fraction of local peaks being also preserved (Figure 5B, S5C). In contrast, GapRMUT was excluded from the mitotic chromatin and remained as low as in interphase (Figure 5A, B). To quantify the increase in GapR binding outside of its localized peaks in interphase, we compared GapR ChIP reads mapping in gene desert regions (> 1 Mb) that lack peaks in asynchronous cells. We detected a 2.5-fold increase in GapR signal during mitosis compared to asynchronous cells (Figure S5D, E), a significant increase that was validated by GapR ChIP-qPCR at specific gene desert loci (Figure S5F, G).

**Fig. 5.**
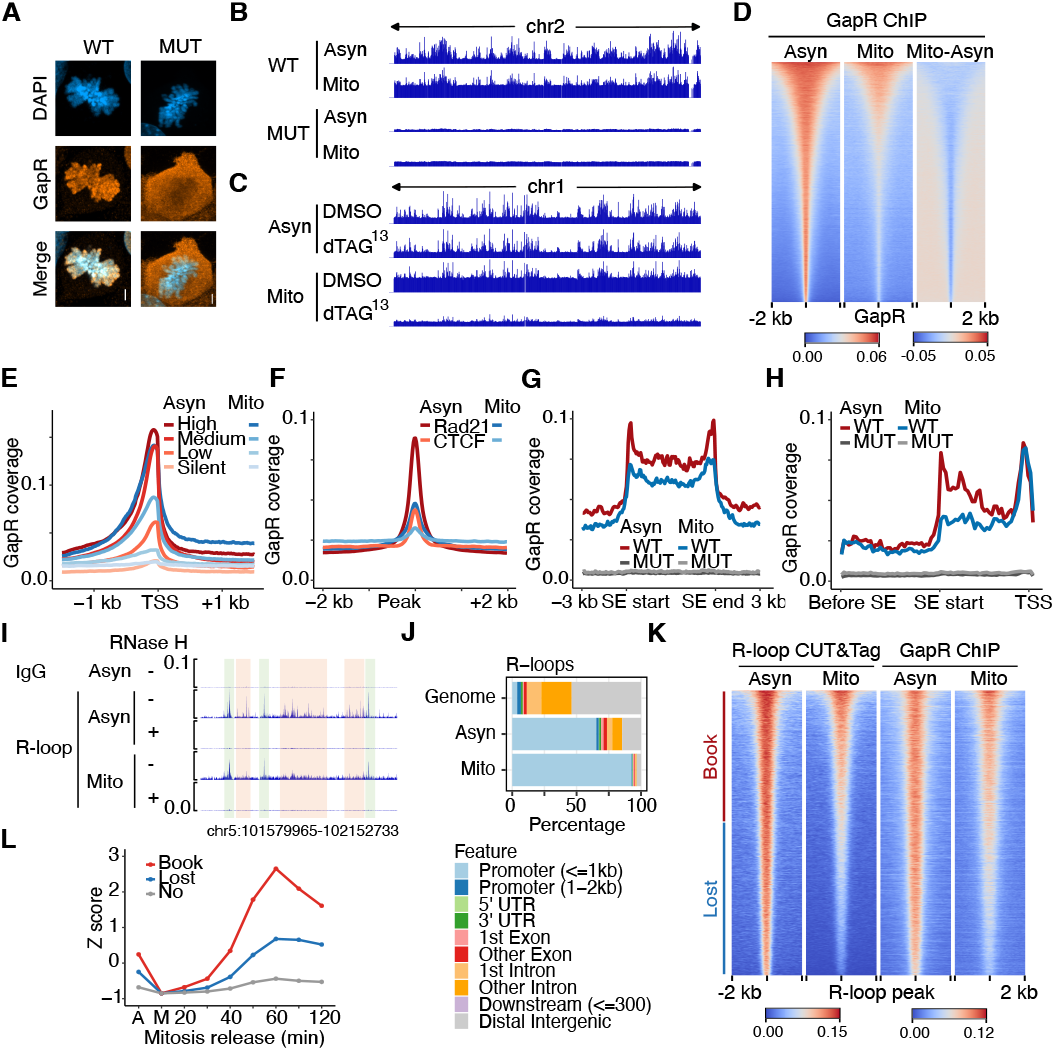
A wave of Condensin driven global positive supercoiling in mitotic cells and the preservation of R-loops at selected promoters prone for rapid post-mitotic transcription. (A) Representative immunofluorescence images of mitotic cells expressing GapRWT/MUT stained with DAPI and anti-FLAG antibody. Scale bar represents 5 µm. (B) Illustrative coverage of E. coli spike-in normalized GapRWT/MUT-FLAG ChIP signal across an entire chromosome in asynchronous (Asyn) and mitotic cells (Mito). (C) Illustrative coverage of E. coli spike-in normalized GapRWT-FLAG ChIP signal across an entire chromosome in asynchronous and mitotic cells treated with DMSO or dTAG13 for 1 h. Mitotic cells were permanently kept with nocodazole. (D). Average heatmaps (n=3) of GapR peaks showing ChIP signal in asynchronous and mitotic cells, ranked by mitotic signal. (E) Average profiles of GapR signal at TSS in asynchronous and mitotic cells across the four Pol II categories shown in Figure 1B. (F) Average profiles of GapR signal at Rad21-CTCF (Rad21) and CTCF only sites (CTCF) in asynchronous and mitotic cells expressing GapRWT/MUT. (G) Average profiles of GapR signal at SEs in asynchronous and mitotic cells expressing GapRWT/MUT. (H) Average binned profiles of GapR signal at SEs and their linked genes. (I) Illustrative coverage of MCF-7 spike-in normalized R-loop CUTTag in asynchronous and mitotic cells. As a control, R-loops were depleted by treating bead-bound cells with RNase H for 2 h. (J) Enrichment barplots showing gene-focused annotations of R-loop CUTTag peaks relative to the genome. (K) Average heatmaps of R-loops showing R-loop CUTTag and GapR signal sorted by mitotic R-loop CUTTag signal. (L) Average post-mitotic pre-mRNA dynamics for genes with retained (Book) or Lost R-loop peaks in mitosis. The “No” category represents genes lacking R-loop CUTTag peaks.

We next asked which factor drives the global increase in positive supercoiling during mitosis. Condensins were previously shown to introduce positive supercoiling into plasmid DNA in vitro and in budding yeast (Kimura and Hirano 1997; Baxter et al. 2011). To assess their role in the mammalian genome, we tagged Smc2, the core subunit of Condensin complexes, with dTAG in ES cells expressing GapR (Figure S5H). Smc2-dTAG was rapidly degraded upon induction (Figure S5I) and in the absence of degradation, ES cells exhibited normal growth, retained normal self-renewing capacity, and did not alter GapR expression (Figure S5J-L).

Thus, we isolated prometaphase arrested cells using nocodazole and depleted Smc2 for 1 h in the continued presence of nocodazole, with all cells remaining mitotic (Figure S5M). As a control, we depleted Smc2 in asynchronous cells for 1h. Consistent with our hypothesis, Smc2 depletion for 1 h in mitotic cells led to a more than four-fold reduction in global GapR binding (Figure 5C, S5N), suggesting that Condensins trigger global positive supercoiling during mitosis. Smc2-depleted asynchronous cells also showed a minor reduction in global GapR signal (Figure 5C, S5N), in line with a role of Condensins in chromatin compaction during interphase in ES cells (Fazzio and Panning 2010).

For the first time, therefore, the hypothesized increase of positive supercoiling during mitosis driven by Condensins is experimentally confirmed and documented both by imaging and ChIP-seq.

### Selected regulatory elements maintain elevated positive supercoiling and R-loops in mitosis

Despite the global increase, clear peaks were nevertheless maintained in mitotic cells (Figure 5B). Indeed, around 25% of peaks identified in interphase were still robustly detected in mitosis (Figure S5C). To clarify whether the loss of 75% of peaks was due to higher global levels, we specifically quantified all GapR peaks. The large majority showed a clear reduction, indicating that, in general, the local mechanisms driving positive supercoiling in interphase are not any longer operational in mitosis (Figure 5D). At the same time, this analysis underscored that the remaining mitotic peaks represent true local activities acting in mitosis, surpassing the global enrichment that takes place. Notably, the remaining peaks in mitosis were highly enriched near promoters compared to asynchronous cells, and were depleted at bivalent, enhancers and insulator regions (Figure S6A). Accordingly, average profiles of distinct regulatory regions illustrate preserved GapR signal at promoters (Figure 5E), but not at enhancers (Figure S6B) or at TAD boundaries (Figure 5F). Interestingly, SEs still retained GapR signal during mitosis (Figure 5G), but the directional component observed in asynchronous cells was lost (Figure 5H). This likely reflects loss of strong and specific enhancer-promoter contacts in mitosis. Like in interphase, we also ruled out that the focal enrichment for GapR in mitosis did not reflect a mere bias to accessibility, given that unique peaks to both GapR (8k) and ATAC (7.5k) can be identified (Figure S6C).

Since transcription and Cohesin-mediated loop extrusion are strongly decreased during mitosis (Gibcus et al. 2018; Ito and Zaret 2022), we turned our attention to Condensins and R-loops. When we quantified positive supercoiling at TSS, TES, enhancers and SEs upon Smc2 depletion, we observed that the effects in mitosis were much stronger than in asynchronous cells (Figure S6D-I), reflecting that the global increase in positive supercoiling takes over at regulatory elements too. However, even in the absence of Condensins activity, clear peaks were still visible in mitosis (Figure S6J). Thus, we aimed at addressing whether these remaining peaks would still be associated with R-loops. To test this, we performed R-loop CUTTag in mitotic cells, using MCF-7 cells as a spike-in control. Consistent with the global shutdown of transcription during mitosis, we observed a significant decrease in the number of R-loop peaks, from 17k in asynchronous cells to 7k in mitosis (Figure 5I, S6K). The mitotic peaks were almost exclusively present near promoter regions (Figure 5J) and, overall, they all showed reduced signal in mitosis. Nevertheless, residual R-loop signal in mitosis quantitatively correlated with GapR signal (Figure 5K), suggesting that these remaining R-loops contribute to local positive supercoiling during mitosis.

The mitotic presence of R-loops and focal positive supercoiling at selected loci, mainly promoters and SEs but also some enhancers (Figure 5J, S6A), is reminiscent of mitotic bookmarking processes, whereby key regulatory elements are differentially regulated during mitosis to promote their fast reactivation upon re-entry into interphase, thereby preserving cell identity. Supporting this, the top mitotic GapR peaks near enhancers and TSSs were associated with mechanisms related to pluripotency (Figure S6L, M). To test this more directly, we classified the R-loop mitotic peaks enriched in positive supercoiling as “bookmarked” and the rest as “lost” (Figure 5K) and compared the transcription dynamics of their associated genes during the transition from mitosis to G1 (Chervova et al. 2023). Interestingly, the book-marked genes showed faster and strongest post-mitotic reactivation than lost genes (Figure 5L, S6N). This suggests that R-loops and the associated positive supercoiling may contribute to the faithful re-establishment of transcription following the mitotic shutdown, suggesting a new role for these structures as drivers of mitotic memory.

Taken together, our results demonstrate that the mitotic genome is globally positively supercoiled. Locally, positive supercoiling and its associated R-loops are retained near TSSs and SEs, potentially serving as a new epigenetic mechanism based on a topological memory that triggers accurate genome reactivation after mitosis.

## Discussion

In this study, we present a comprehensive and functional map of positive DNA supercoiling during interphase and mitosis in mouse ES cells. Through the dissection of the mechanisms driving their biogenesis, we have identified several features with profound mechanistic implications. First, our findings align with the twin-domain model of supercoiling, revealing localized accumulation of positive supercoiling immediately adjacent to negatively supercoiled regions such as R-loops, especially at promoters, as well as broad domains of positive supercoiling downstream of TESs, spanning kilobase-long regions of DNA. Secondly, and in contrast to prevailing views, we show that positive supercoiling accumulates at active gene regulatory elements critical for transcription, 3D organization, and maintenance of cell identity. Notably, while Topoisomerases resolve positive supercoiling genome-wide, we show a high diversity of molecular sources, with transcription being critical at gene ends, R-loops at promoters, enhancers and super-enhancers, and Cohesin at loop anchors involved in TADs formation and enhancer-promoter interactions. Thirdly, we provide evidence for a long-standing hypothesis of Condensin-dependent positive supercoiling in mitosis. Finally, we identify persistent local topological features in mitotic chromatin characterized by Condensins and R-loops driven positive supercoiling, especially at promoters that are rapidly reactivated after mitosis. Below, we discuss these features in regard to their potential functions in genome regulation.

Identifying positive supercoiling at promoters and linking it to the presence of R-loops rather than to transcriptional activity itself was counter-intuitive, even though this has been observed in yeast and in human cell lines (Guo et al. 2021; Longo et al. 2024). A priori, positive supercoiling would have a negative impact on transcription, since over wounded DNA may inhibit transcription factor binding and DNA melting by the transcriptional machinery. In the meantime, R-loops have opposing effects on transcription, stabilizing DNA opening but inducing Pol II pausing (Belotserkovskii et al. 2017; Chen et al. 2017). Indeed, Topoisomerases are needed to resolve R-loops and enable productive Pol II elongation. In this regard, it is possible that the mutually reinforcing nature of R-loops and adjacent positive supercoiling may increase the recruitment of Topoisomerases to finely tune transcriptional bursting (Jonkers and Lis 2015; Chen et al. 2017; Hwang et al. 2025). In such a model, R-loops and positive supercoiling would form a dynamic, reversible and cyclic regulatory phenomenon orchestrated by Topoisomerases on timescales relevant to transcriptional bursting (Ancona et al. 2019; Hwang et al. 2025).

Finding positive supercoiling at both extremities of active genes suggests that transcribed units might be viewed as partially self-contained, topologically insulated units, optimizing transcription bursting, R-loop formation, nucleosome dynamics, binding of transcription factors and Pol II recycling, while limiting excessive interference within neighboring regions (Perales and Bentley 2009; Ancona et al. 2019; Fosado et al. 2021). In this regard, the enhancement of positive supercoiling at promoters and 3’ ends mediated by Cohesin, most likely through its DNA extrusion activity, suggests a new role for this molecular motor in topologically insulating active genes by alternative mechanisms to proper 3D loops.

Notably, both the impact of R-loops and Cohesin is not limited to promoters, but extends to virtually all classes of regulatory elements, especially at enhancers and superenhancers. Similar to the model proposed for promoters, it is tempting to speculate that the dynamic activity generated by R-loops, positive-supercoiling and Topoisomerases may also lead to a fine-tunning of enhancer function characterized by alternate periods of competence to bind transcription factors and refractory periods where positive supercoiling takes over. This functional breathing of enhancers might be mechanistically related to their pulsatile activity and their influence of transcriptional bursting (Fukaya 2023). Thus, positive supercoiling at gene regulatory elements may be instrumental to finally tune the temporal discreet dynamics of enhancer and promoter activity.

In addition to separately influence enhancer and promoter activity, positive supercoiling may also impact how enhancers directly control promoters. Indeed, positive supercoiling also characterizes all loop anchors, whether forming TADs or enhancer-promoter and promoter-promoter loops, and, strikingly, the intervening regions. Since Cohesin depletion strongly affects positive supercoiling within loops, as suggested by in vitro and modelling approaches (Sun et al. 2013; Racko et al. 2018), it seems likely that it directly contributes to their formation or stabilization, fostering contacts between distal regions, especially in the context of directional super-enhancer contacts with their target promoters. The observation that positive supercoiling and its dependency to Cohesins scales with Cohesin and CTCF binding at loop anchors further provides an additional model to explain differences in the insulating capacity of distinct CTCF binding sites (Papale et al. 2025), which may provide a local barrier to the spreading of topological effects in a way analogous to the model proposed for transcriptional units.

In addition to active gene regulatory processes, the induction of positive supercoiling by loop extrusion appears a general property of this class of DNA motors, since it is not specific to Cohesin but also applies to Condensins (Kimura and Hirano 1997; Bazett-Jones et al. 2002). Documenting for the first time the increase of positive supercoiling driven by Condensins, the main players in chromatin organization during mitosis, provides new explanations to the specificities of mitotic chromatin. Notably, it is noteworthy that the 2.5 fold global increase in positive supercoiling occurring during mitosis correlates extremely well with the 2-3 fold increase in chromatin compaction measured independently (Paulson et al. 2021). This suggests that the global wave of positive supercoiling mediated by Condensins actively contributes to the compaction of mitotic chromosomes, beyond their active role in dissolving TADs and restructuring the 3D folding of the chromatin as dynamic, nested loops (Gibcus et al. 2018). Moreover, this global excess of positive supercoiling might contribute to the global loss of transcription factor binding and transcriptional activity during mitosis (Ito and Zaret 2022). However, positive supercoiling may have additional gene regulatory functions. Indeed, a subset of promoters maintain higher positive supercoiling than average and detectable R-loops. In agreement with the seminal proposal of mitotic bookmarking, where the involvement of DNA supercoiling was anticipated (Michelotti et al. 1997), these promoters are strongly and rapidly activated upon mitotic exit, suggesting that residual R-loops and increased super-coiling prime genes for post-mitotic reactivation. Mechanistically, this could be related to a strong local attraction of Topoisomerases during mitotic chromosome decondensation, which would in turn promote transcriptional activity.

Overall, this study suggests that positive supercoiling may have previously unanticipated roles in the fine regulation of transcriptional dynamics, influencing the activity of promoters, enhancers and their 3D contacts, as well as contributing to the establishment of different modes of genome architecture in interphase (TADs) and mitosis (nested loops). Its role as a topological memory of active promoters is, moreover, an exciting possibility to directly link DNA topology with epigenetic information and the perpetuation of gene regulatory networks during mitosis. Future studies will need to design specific experimental strategies to dissect each and all of these new modes of gene regulation.

## Supplementary information

Six supplementary figures accompany this manuscript, they can be found online.

## Acknowledgements

The authors thank Monica S. Guo and Michael T. Laub for providing the WT and mutant GapR plasmids and Carla Saleh’s lab for Drosophila S2 cells. The authors acknowledge the Biomics core facility for access to the NextSeq instruments and Flow Cytometry platform. The work was supported by grants to P.N. from the European Research Council (ERC-CoG-2017 BIND), Agence Nationale de la Recherche (Nucleoseq S-CR22069) and recurrent funding from Institut Pasteur, Revive (Investissement d’Avenir, ANR-10-LABX-73) and the CNRS.

## Author contributions

A.K.S. performed all experiments and bioinformatic analysis with input from L.A-P and A.D. L.T. performed RNA-seq analysis. A.K.S. and P.N. conceived the project, interpreted the results and wrote the manuscript.

