## Supplementary and Methods for "Topological Regulation of the Mammalian Genome by Positive DNA Supercoiling"

### SUPPLEMENTARY FIGURE AND LEGENDS

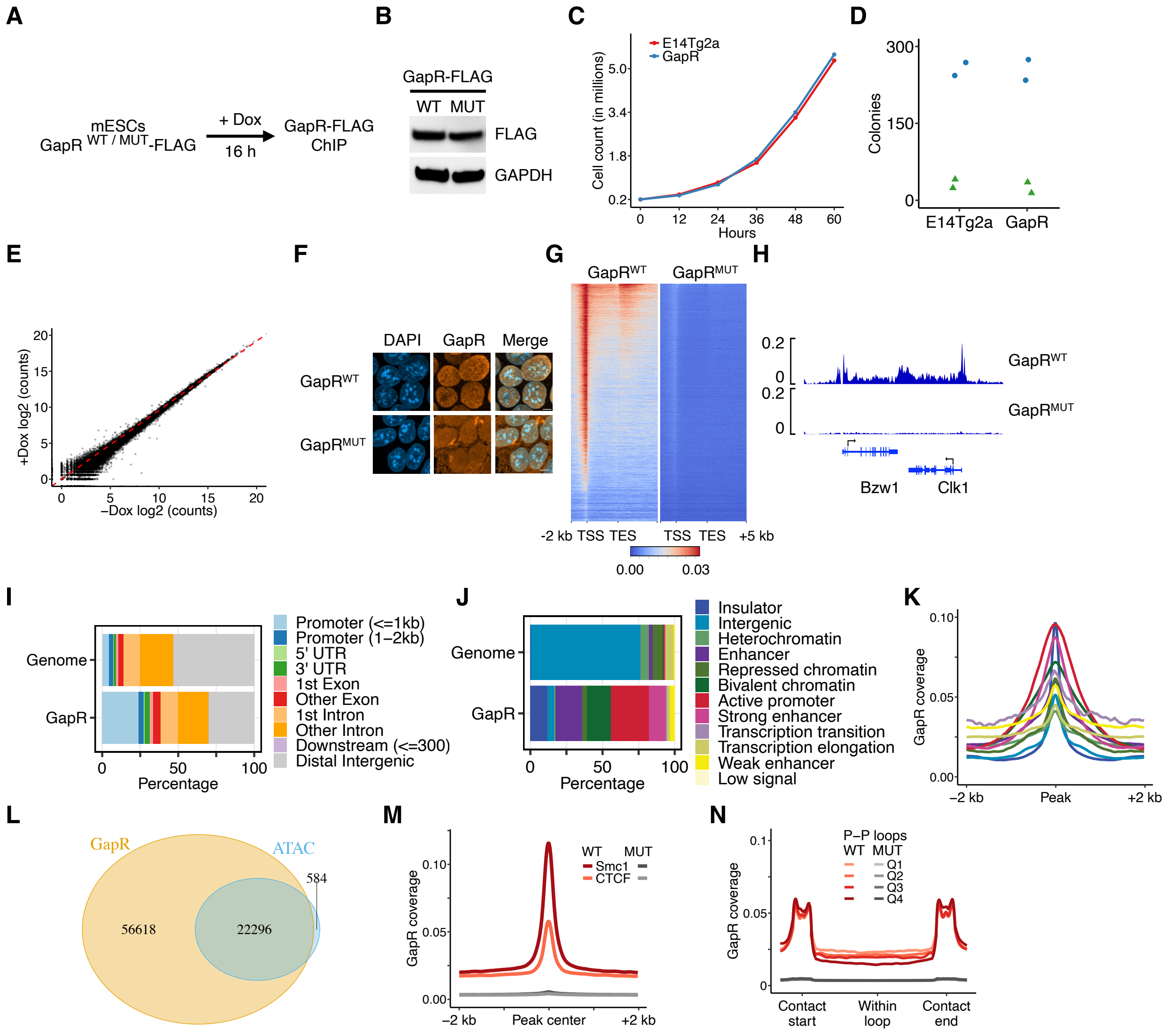


**Figure S1. GapR binding at regulatory elements. (A)** Experimental scheme for GapR-FLAG ChIP following 16 h dox induction in mouse ES cells. **(B)** Western blot showing GapR$^{WT/MUT}$-FLAG and GAPDH expression upon dox induction. **(C)** Growth curve of control (E14Tg2a) and GapR expressing cells induced with dox for 16 h and subsequently cultured in media lacking dox. **(D)** Number of alkaline phosphatase positive (circles) and negative (triangles) colonies from cells described in B. **(E)** RNA-seq of uninduced and dox induced GapR cells for 16 h. **(F)** Representative immunofluorescence images of cells expressing C-terminal FLAG-tagged GapR$^{WT/MUT}$ stained with DAPI and anti-FLAG antibody. Scale bar represents 5 μm. **(G)** Average binned heatmaps of all genes, showing $GapR^{WT}$ and $GapR^{MUT}$ signal sorted by RNA-seq TPM counts. **(H)** Illustrative coverage of spike-in normalized GapR signal between two expressed convergent genes. **(I)** Enrichment barplots showing gene-focused annotations of GapR peaks relative to the genome. **(J)** Bar plot showing enrichment of GapR peaks across ChromHMM categories, relative to the genome. **(K)** Average profiles showing GapR enrichment across ChromHMM regions. **(L)** Venn diagram showing overlap between GapR and ATAC-seq peaks. **(M)** Average profiles of $GapR^{WT/MUT}$ signal centered at Smc1–CTCF co-bound peaks (Smc1) and CTCF-only sites (CTCF). **(N)** Average binned profiles of $GapR^{WT}$ within promoter-promoter (P-P) DNA loops, aligned to the start and end of interacting loop anchors and divided into quartiles based on the loop size. Median loop sizes for each quartile were Q1 = 59 kb, Q2 = 133 kb, Q3 = 250 kb and Q4 = 293 kb.

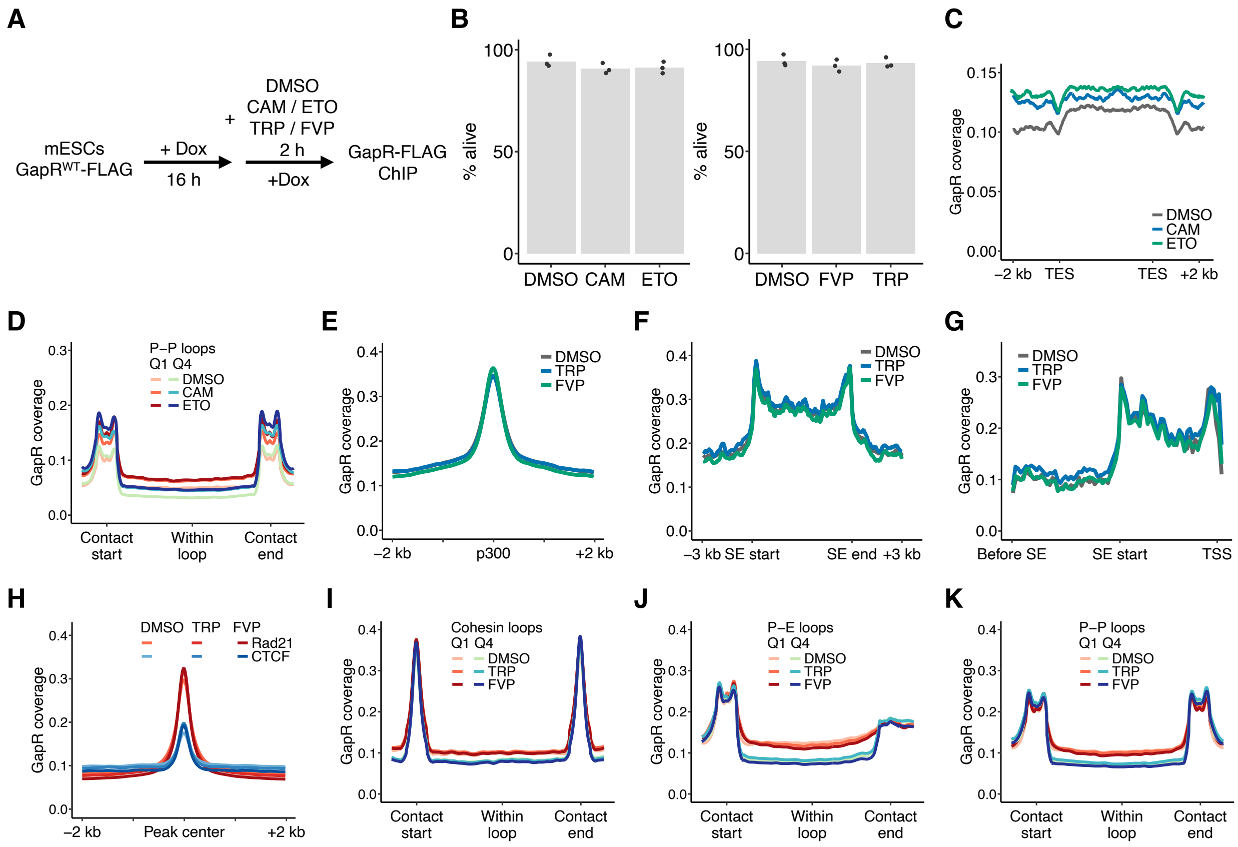


**Figure S2. Effects of topoisomerases and transcription on regulatory elements. (A)** Experimental scheme for GapR$^{WT}$-FLAG ChIP following 16 h dox induction in ES cells and treated for 2 h with DMSO, topoisomerase (CAM and ETO) or transcription inhibitors (TRP and FVP) in the continued presence of dox. **(B)** Barplots showing the percentage of viable cells measured by trypan blue staining after 2 h treatment with DMSO or inhibitors. **(C)** Average binned profiles of GapR signal between expressed convergent genes treated with topoisomerase inhibitors. **(D)** Average binned profiles of GapR within the smallest (Q1) and largest (Q4) P-P loops following topoisomerase inhibition. **(E)** Average profiles of GapR at active enhancers centered on p300 summit. **(F)** Average binned profile of GapR at SEs. **(G)** Average binned profiles of GapR centered on the SE start and spanning an equidistant distant with its target gene and the upstream region. **(H)** Average profiles of GapR signal at Rad21-CTCF (Rad21) and CTCF only sites (CTCF) following transcription inhibition. **(I-K)** Average binned profiles of GapR within the smallest (Q1) and largest (Q4) cohesin (I), E-P (J) and P-P (K) loops following transcription inhibition.

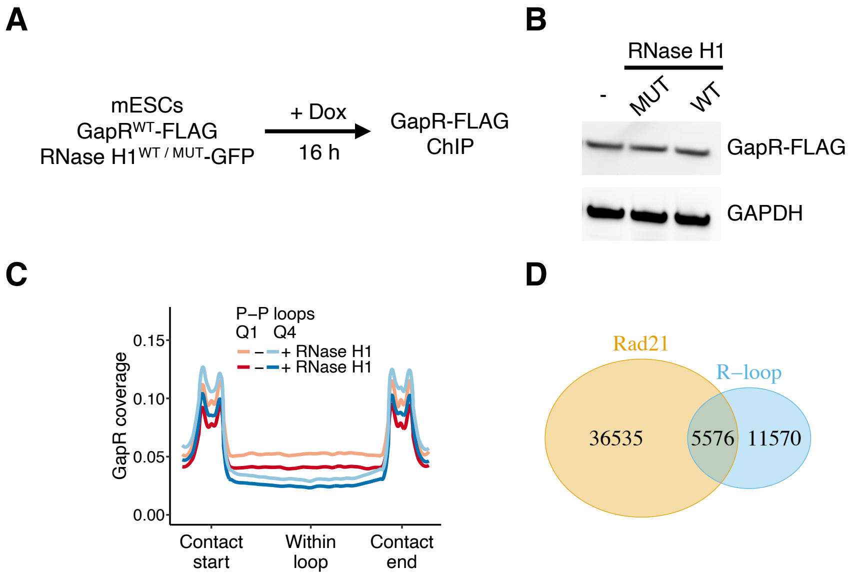


**Figure S3. Effects of RNase H1 expression on regulatory elements. (A)** Experimental scheme for GapR$^{WT}$-FLAG ChIP following 16 h dox induction in ES cells expressing RNase H1$^{WT/MUT}$. **(B)** Western blot showing GapR and GAPDH expression in cells lacking RNase H1 or expressing RNase H1 WT or MUT following 16 h dox induction. **(C)** Average binned profiles of GapR within the smallest (Q1) and largest (Q4) P-P loops following RNase H1 expression. **(D)** Venn diagram showing overlap of Rad21 ChIP and R-loop CUT&Tag peaks.

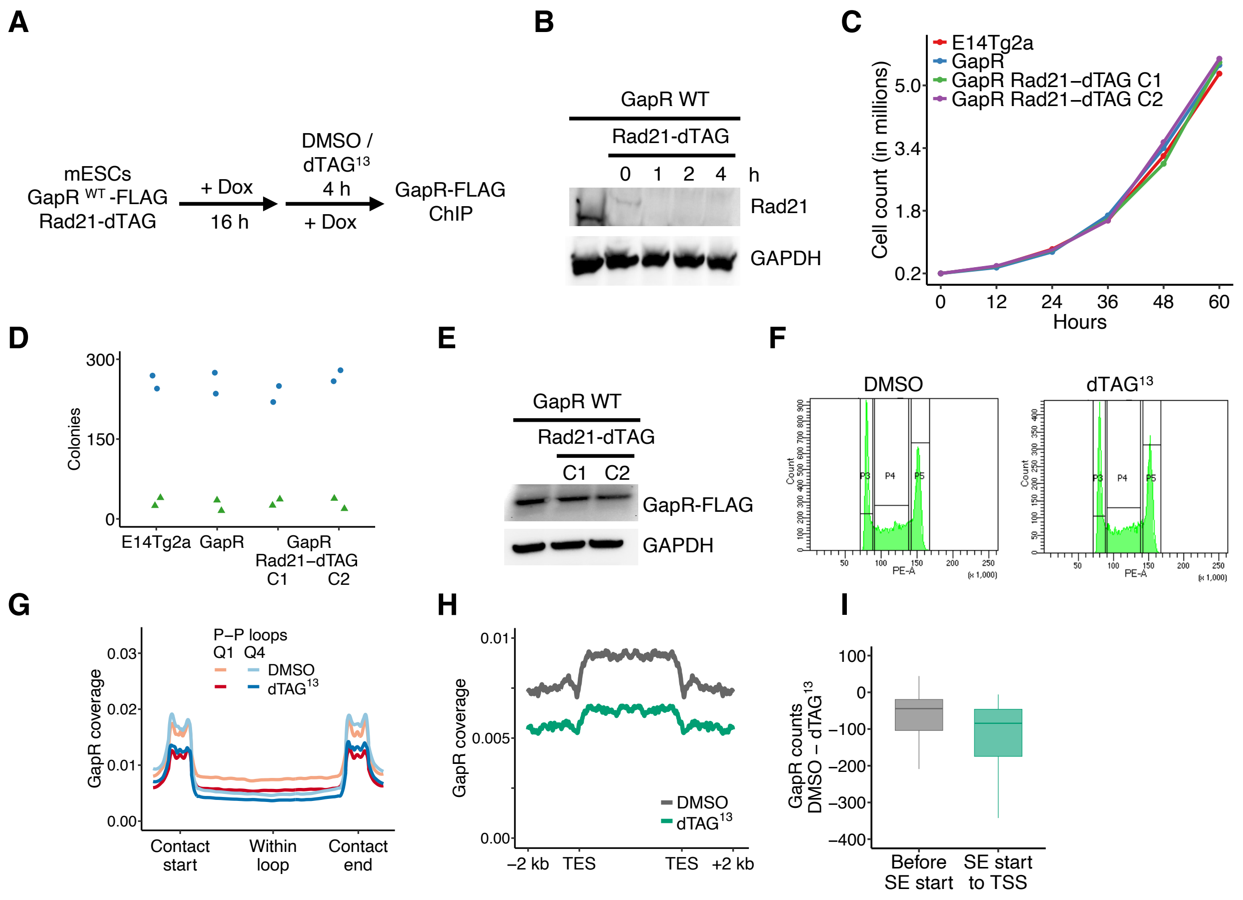


**Figure S4. Effects of Cohesin depletion on regulatory elements. (A)** Experimental scheme for GapR^WT-FLAG ChIP in cells homozygously expressing Rad21- dTAG. **(B)** Western blot showing Rad21-dTAG and GAPDH expression upon dTAG^13 treatment for 0 to 4 h. **(C)** Growth curve of control (E14Tg2a) and GapR expressing cells with or without Rad21-dTAG, induced with dox for 16 h and subsequently cultured without dox. **(D)** Number of alkaline phosphatase positive (circles) and negative (triangles) colonies from cells described in C. **(E)** Western blot showing GapR and GAPDH expression Rad21-dTAG cells in two clones (C1 and C2). **(F)** Cell cycle profiles of Rad21-dTAG cells following DMSO or dTAG^13 treatment for 4 h. **(G)** Average binned profiles of GapR signal within the smallest (Q1) and largest (Q4) P-P loops. **(H)** Average binned profiles of GapR signal between expressed convergent genes. **(I)** Boxplots showing quantification of GapR signal in the region between SE start and TSS and the equidistant region upstream of the SE start. Number of reads overlapping each region were calculated and the difference between DMSO and dTAG^13 treated cells are shown.

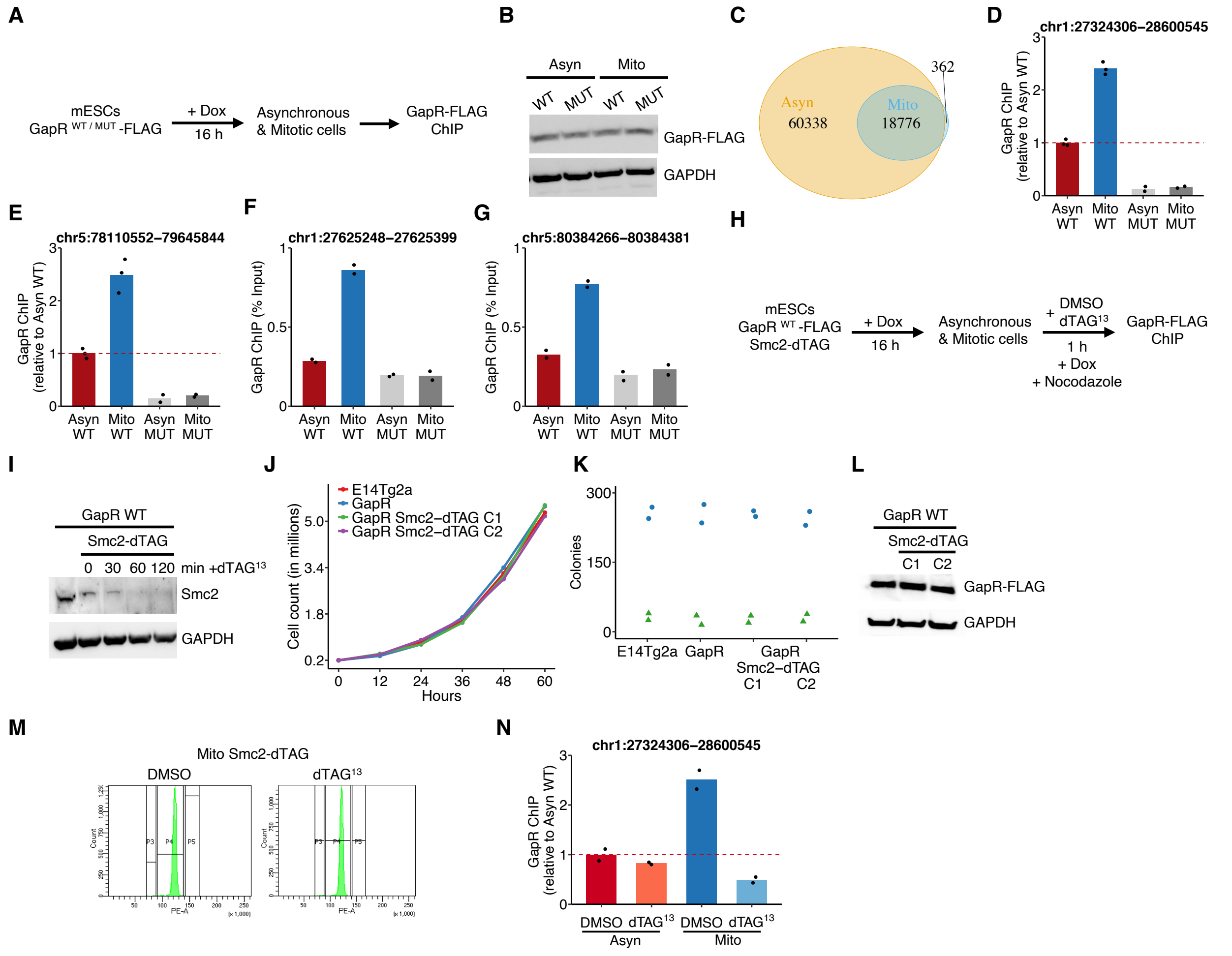


**Figure S5. Characterization of Condensins driven positive supercoiling in mitosis. (A)** Experimental scheme for GapR$^{WT/MUT}$-FLAG ChIP in ES cells treated with dox for 16 h. Mitotic cells (>95% purity) were isolated following nocodazole treatment for 4 h in the continued presence of dox. **(B)** Western blot showing GapR and GAPDH expression in asynchronous and mitotic cells expressing GapR$^{WT/MUT}$. **(C)** Venn diagram showing overlap of GapR peaks in asynchronous and mitotic cells. **(D, E)** Barplots showing relative quantification (Asyn WT set to 1; n=3) of GapR signal in the indicated large regions (>1Mb) lacking GapR peaks. **(F, G)** Barplots showing ChIP-qPCR quantification of GapR binding at the indicated loci (mm10) in asynchronous and mitotic cells expressing GapR$^{WT/MUT}$. **(H)** Experimental scheme for GapR$^{WT}$-FLAG ChIP in cells homozygously expressing Smc2-dTAG, treated with DMSO or dTAG$^{13}$ for 1 h in the continued presence of dox. **(I)** Western blot showing Smc2-dTAG and GAPDH expression upon treatment with dTAG$^{13}$ in mitotic cells. **(J)** Growth curve of control (E14Tg2a)

and GapR expressing cells with or without Smc2-dTAG construct, induced with dox for 16 h and subsequently cultured in media lacking dox. **(K)** Number of alkaline phosphatase positive (circles) and negative (triangles) colonies from cells described in J. **(L)** Western blot showing GapR and GAPDH expression in cells harboring Smc2-dTAG construct. C1 and C2 denote two clones used for GapR ChIP-seq. **(M)** Cell cycle profiles of Smc2-dTAG mitotic cells following DMSO or $dTAG^{13}$ treatment for 1 h. **(N)** Barplots showing relative quantification (Asyn WT set to 1; n=2) of GapR signal in the indicated large region (>1Mb) lacking GapR peaks.

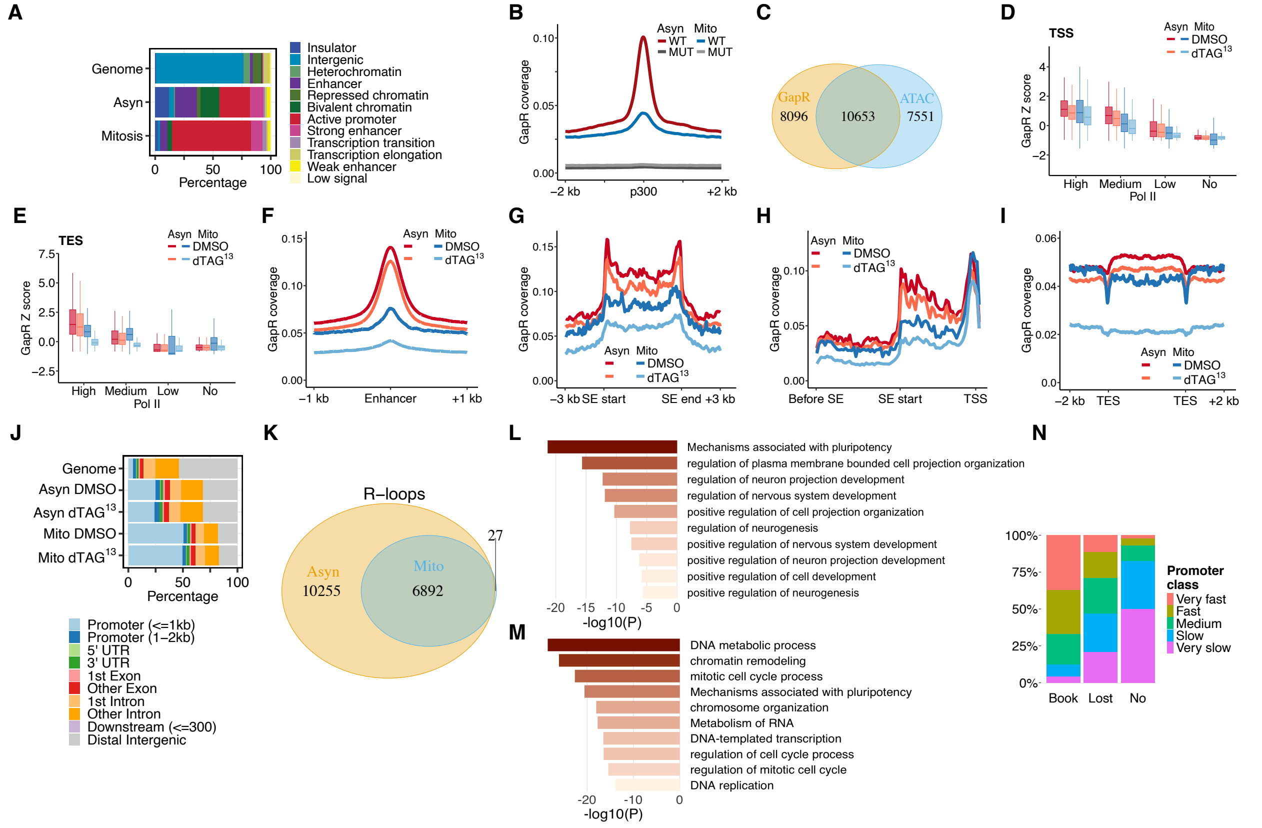


**Figure S6. Effects of GapR and R-loops near gene regulatory elements in mitosis. (A)** Bar plot showing enrichment of GapR peaks across ChromHMM categories in asynchronous and mitotic cells, relative to the genome. **(B)** Average profile of GapR signal at active enhancers centered on p300 summits in asynchronous and mitotic cells expressing $GapR^{WT/MUT}$. **(C)** Venn diagram showing overlap of GapR and ATAC-seq peaks in mitotic cells. **(D)** Boxplots quantifying GapR signal (n=2) from TSS across the four Pol II categories using the [-1, 0.2 kb] interval in asynchronous and mitotic cells Smc2-dTAG treated with DMSO or $dTAG^{13}$. **(E)** Same as D, but for signal downstream of TES (5kb). **(F)** Average profiles of GapR ChIP signal at active enhancers centered on p300 summits in asynchronous and mitotic cells treated with DMSO or $dTAG^{13}$. **(G-I)** Average binned profiles of GapR signal at SEs (G), at the regions liking them with their target genes (H) or at convergent genes (I), in asynchronous and mitotic cells treated with DMSO or $dTAG^{13}$. **(J)** Enrichment bar plots showing distribution of gene-focused annotations for GapR peaks in asynchronous and mitotic Smc2-dTAG cells treated with DMSO or dTAG[13]. **(K)** Venn diagram showing overlap of R-loop CUT&Tag peaks in Asyn and Mito cells. **(L, M)** Gene ontology terms associated with the nearest genes of the top 2000 GapR mitotic peaks overlapping p300 sites in interphase (L) or of the genes driven by promoters overlapped by top 2000 mitotic GapR peaks overlapping TSSs (M). **(N)** Stacked barplots showing the percentage of genes displaying different post-mitotic transcriptional dynamics, as previously established (Chervova et al. 2023), as a function of the overlap of their promoters with R-loops peaks characterized as book, lost or absent.

### METHODS

#### Cell culture

*General:* Mouse embryonic stem cells (E14Tg2a) were grown in DMEM + GlutaMax-I (Gibco, Cat#31966-021) supplemented with 10% FBS (Sigma, Cat#F7524), 100 μM β-mercaptoethanol (Gibco, Cat#31350-010), 1X MEM non-essential amino acids (Gibco, Cat#1140-035) and 10 ng/ml recombinant mouse LIF (Miltenyi Biotec, Cat#130-099-895) on 0.1% Gelatin (Sigma, Cat#G1890-100G) at 37 °C with 7% $CO_2$. For passaging, cells were washed with phosphate-buffered saline (PBS) and dissociated with 1X Trypsin-EDTA for 2-3 min (Gibco, Cat#25300-054). Cells were diluted 1:5 in fresh media and transferred into new culture falcons. Cells were regularly tested for mycoplasma contamination (Biovalley, Cat#11-8100).

Michigan Cancer Foundation-7 (MCF-7) cells, used as a spike in for CUT&Tag experiments, were cultured under the same conditions as mouse ES cells, except without gelatin, LIF and passaged with 1:3 dilution. *Drosophila* Schneider Line 2 (S2) cells, used as a spike in for RNA-seq experiments, were maintained at 28 °C in Schneider's drosophila medium (Gibco Cat#21720024) with 10% FCS and penicillin-streptomycin 1:200 dilution (Gibco, Cat#15140148).

*Preparation of mitotic cells:* ES cells were cultured in T150 flasks (Dutscher, Cat#190150) to 70-80% confluency. Cells were vigorously tapped to remove loosely attached colonies and dead cells. Medium was discarded and cells were incubated in fresh medium with 50 ng/ml nocodazole (Sigma, Cat#487929) for 4 h. Cells were washed with once with PBS and gently tapped in 10 ml medium. The cell suspension was filtered through 10 μm filter (pluriSelect, Cat#43-50010-03) and collected by centrifugation. Preparations with >95% purity were used for further experiments.

*Clonal assay:* For each cell line, 600 cells were plated per well in a 6 well culture plate (TPP, Cat#92006), with three technical replicates, in standard LIF medium. Medium was changed daily for 6 days. Colonies were washed once with PBS, fixed with 1 ml fixing solution per well (25% citrate solution, 67% acetone, 8% formaldehyde) for 1 min. Colonies were washed with water and stained with 1 ml staining solution (Alkaline phosphatase staining kit, Sigma, Cat#86R-1KT) for 20 min. Colonies were washed with water and air-dried overnight. Undifferentiated and differentiated colonies were counted and averaged.

*Growth assay:* 0.2 x $10^6$ cells were seeded per well in a 6 well culture plate, harvested from a well every 12 h with trypsin and counted in triplicate.

**Generation of cell lines**

*GapR expressing ES cells:* GapR WT and MUT (amino acids 1-76) genes with C-terminal 3xFLAG tag were amplified from pKVS45-GapR-3xFLAG and pKVS45-gapR(1-76)-3xFLAG plasmids (Guo et al. 2021), cloned into a plasmid with TetO, β-globin polyA and a puromycin selection cassette (Festuccia et al. 2016) and verified by Sanger sequencing. Linearized plasmids were electroporated in E14Tg2a cells having a rTta-M2 construct. After selection with puromycin 1 µg/ml (Sigma, Cat#P8833) for 6-7 days, individual clones were expanded and induced with 1 µg/ml dox overnight to screen GapR-FLAG expression by Western-Blot. Clones with highest and similar $GapR^{WT/MUT}$ expression were selected. For most experiments, cells were induced 1 µg/ml dox overnight, washed with PBS and collected with standard trypsinization.

*RNase $H1^{WT/MUT}$ in $GapR^{WT}$ ES cells:* E14Tg2a cells stably expressing $GapR^{WT}$ were used to generate lines expressing RNase $H1^{WT/MUT}$ under dox control. The pEGFP-N2-2xNLS-RNase $H1^{WT/MUT}$ constructs were PCR amplified from plasmids (Addgene Cat#196702 and 196703) and cloned as described above but with a Neomycin selection cassette. Linearized plasmids were electroporated into GapR expressing cells, which were then selected with G418 350 µg/ml (Roche, Cat#4727878001) for 6-7 days. Individual clones were expanded and induced with 5 µg/ml dox overnight to co-induce both GapR and RNase H1. RNase H1 expression was tested by eGFP expression in live cells and for GapR-FLAG expression with western blot. First, clones with highest eGFP expression were selected. Then clones with similar GapR expression were chosen. For ChIP-seq experiments, control cells without RNase H1 construct and cells harboring RNase $H1^{WT/MUT}$ constructs were induced with 5 µg/ml dox overnight, washed with PBS and collected with standard trypsinization.

*Rad21-dTAG and Smc2-dTAG in GapR$^{WT}$ ES cells:* E14Tg2a cells expressing GapR$^{WT}$ were used to generate Rad21 and Smc2 degron lines with the FKBP12$^{F36V}$ variant. Guide RNA (gRNA) oligos targeting the C-terminus of Rad21 (CACCGCCACGGTTCCATATTATCTG and AAACCAGATAATATGGAACCGTGGC) and Smc2 (CACCGAACTGGCATGCACTCAGTT and AAACAACTGAGTGCATGCCAGTTC) were synthesized from Integrated DNA Technologies, annealed in NEB Buffer 2 (Cat#B7002S) and ligated into a plasmid containing *Sp*Cas9 (Addgene, Cat#64324). Homology arms (~400 bp) flanking the Rad21 stop codon and Smc2 stop were cloned into a plasmid containing a GS linker followed by FKBP12$^{F36V}$-loxP-Hygromycin cassette-loxP. Clones were sequenced using Sanger sequencing and nucleofected into ES cells (5 M) using the mouse ES cell nucleofector kit (Lonza, Cat#VPH-1001) according to the manufacturer's instructions. Cells were recovered for 2 days in standard LIF medium, and then selected for 6-7 days with hygromycin 150 µg/mL (Sigma, Cat#SBR00039). Individual clones were picked and cultured with dTAG[13] (Sigma, Cat#SML2601) for 2-3 days for initial selection, as Rad21 and Smc2 are essential for cell viability. At least two clones were further selected by PCR using genomic DNA at the Rad21 and Smc2 loci.

**Topoisomerase and transcription inhibition:** For each condition, 2 x $10^6$ cells were cultured in T75 flasks under standard conditions with 1 µg/ml dox overnight. Cells were washed with PBS and incubated in fresh medium containing 1 µg/ml dox and either inhibitors or an equal volume of DMSO (Sigma, Cat#D2650) for 2 h. Cells were washed with PBS and collected by standard trypsin dissociation. Inhibitors were used at following concentrations: Triptolide - 1 µM (Sigma, Cat#T3652), Flavopiridol - 1 µM (Sigma, Cat#F3055), Camptothecin - 100 µM (Sigma, Cat#C9911), Etoposide - 100 µM (Sigma, Cat#E1383).

**Rad21 and Smc2 dTAG-13 depletion:** To deplete Rad21 and Smc2 proteins in asynchronous cells expressing GapR, cells were cultured overnight with 1 µg/ml dox. Next day, cells were washed once with PBS and incubated in fresh medium containing 1 µg/ml dox and either 500 µM dTAG[13] or an equal volume of DMSO.

For Smc2 depletion during mitosis, cells were cultured overnight with 1 µg/ml dox to induce GapR expression. Mitotic cells were prepared as above, with all washing and incubation steps included 1 µg/ml dox to maintain GapR expression and 50 ng/ml nocodazole to prevent mitotic exit. To induce Smc2 depletion, cells were diluted to 1 x $10^6$/ml in fresh media with 1 µg/ml dox and 50 ng/ml nocodazole and incubated with either 500 µM dTAG[13] or an equal volume of DMSO for 1 h in a petri dish (Dutscher, Cat#067003). After 1 h depletion, cells were collected by gentle pipetting and centrifugation.

**Western blot:** 0.5 x $10^6$ cells were lysed in 100 µl Laemmli Buffer (Bio-Rad, Cat#1610737) at 95 °C for 5 min. 50 µl was loaded on a precast protein gel (Bio-Rad, Cat#456-1034), migrated at 15 V/cm in MOPS SDS running buffer (Millipore, Cat#MP81G15) and transferred to nitrocellulose membrane using iBlot 2 Transfer Stacks (Invitrogen, Cat#IB23001). Membrane was blocked with 5% BSA (Sigma, Cat#10711454001) in PBS with 0.1% Tween-20 (PBST) and incubated overnight with primary antibodies (anti-FLAG (Sigma, Cat#F1804, 1:10000 dilution), anti-Smc2 (Cell Signaling Technology, Cat#D11F9, 1:1000 dilution), anti-Rad21 (Bethyl Laboratories, Cat#A300-080A, 1:10000 dilution)).

**Imaging:** ES cells were cultured on poly-L-ornithine/laminin treated ibidi plates, fixed with 3.7% formaldehyde and washed twice with PBS. Cells were permeabilized with PBS + 0.5% Triton X-100, washed with PBS, and blocked with PBS supplemented with 3% donkey serum (Sigma, Cat# D9663) for 15 min on ice. Primary antibodies (in PBS + 3% donkey serum) were applied for 2 h overnight at 4°C. Cells were washed thrice with PBS and incubated with secondary antibodies (in PBS with 3% donkey serum) for 2 h at RT. Cells were washed thrice with PBS, nuclei counterstained with DAPI (Sigma, Cat# D9542), and imaged with a confocal microscope. Antibodies used were as follows:

1. Anti-FLAG (Sigma, Cat#F1804), 1:100 dilution
2. Alexa Fluor 488 Donkey Anti-Mouse (Thermo Fisher, Cat#A-21202), 1:500 dilution

**Cell cycle analysis:** Cells were fixed in 70% ethanol at 4 °C for 2 h and resuspended in 500 µl PBS. Cells were incubated with 0.2 µg/µl RNase A for 30 min at 37°C and with 20 µg/ml propidium iodide for 1 h. Cell cycle was analyzed on BD FACSymphony A5 cytometer.

**Spike-in ChIP-seq:** ES cells and MCF-7 cells were crosslinked with 1% formaldehyde (Thermo Fisher, Cat#10751395) at 5 x $10^6$ cells/ml for 10 min at room temperature (RT) on a rotating wheel, followed by 0.125 M glycine for 5 min. Cells were centrifuged (5 min, RT, 300x*g*), washed twice with ice-cold PBS and resuspended at 5 x $10^6$ cells/ml in a swelling buffer (25 mM HEPES pH 8, 10 mM KCl, 10 mM EDTA, 0.5% NP-40, 1x protease inhibitors (Roche, Cat#04693116001)) for 30 min on ice. Cells were homogenized with a douncer for 30 times (for asynchronous cells only), centrifuged (5 min, 4 °C, 600x*g*) and resuspended at $10^7$ cells/ml in TSE150 (0.1% SDS, 1% Triton X-100, 2 mM EDTA pH 8, 20 mM Tris-Cl pH 8, 150 mM NaCl, 1x protease inhibitors) buffer. Nuclei were sonicated in 1.5 ml tubes (Diagenode, Cat#C30010016) in Bioruptor for 7 cycles of 30'' ON/30'' OFF cycles at 4 °C. Chromatin was centrifuged (10 min, 4 °C, 11000x*g*), supernatant was flash frozen and stored at -80 °C. Genomic DNA was isolated from a fraction of chromatin and measured using Qubit fluorometer.

To prepare DNA from *E. coli* expressing $GapR^{WT}$ for spike in, *E. coli* was transformed with pKVS45-GapR-3xFLAG plasmid and grown overnight in LB medium to OD 0.3. Next day, cells were diluted to OD 0.01 and $GapR^{WT}$ expression was induced with 25 ng/mL anhydrotetracycline (Takara Bio, Cat#631310) for 2 h. Cells were fixed with 10 mM sodium phosphate pH 7.6 and 1% formaldehyde for 10 min at RT on a rotating wheel followed by 0.1 M glycine for 5 min. Cells were washed thrice with PBS, resuspended in 500 µl TES buffer (10 mM Tris-Cl pH 7.5, 1 mM EDTA, 100 mM NaCl) and incubated with 35000 U Lysozyme (Bioresearch Technologies, Cat#R1804M) for 15 min at RT followed by addition of 500 µl ChIP buffer (16.7 mM Tris-Cl pH 8, 167 mM NaCl, 1.1% Triton X-100, 1.2 mM EDTA, protease inhibitors). Lysates were incubated at 37 °C for 10 min, sonicated in 1.5 ml Bioruptor tubes for 10 cycles of 30'' ON/30'' OFF cycles at 4 °C and supernatant was recovered by centrifugation (10 min, 4 °C, 11000x*g*). Genomic DNA was isolated from a fraction of supernatant and measured using Qubit fluorometer.

For each ChIP, 90% ES cell chromatin (from 3-5 x $10^6$ cells) and 10% *E. coli* spike were mixed in a 500 µl total volume, mixed for 10 min at 4 °C. 10 µl was saved as an input and rest was incubated with 3 µg anti-FLAG M2 (Sigma, Cat#F1804), anti-Pol II (Santa Cruz, Cat#N-20 and

H-224) antibody overnight at 4 °C. Pre-blocked protein G Sepharose beads (Sigma, Cat#P3296-5 ML) were added and incubated for 4 h. Beads were washed with 1 ml buffer in the following order: thrice with TSE150, once with TSE500 (TSE150 but 500 mM NaCl), once with washing buffer (10 mM Tris-Cl pH 8, 0.25 M LiCl, 0.5% NP-40, 0.5% Na-deoxycholate, 1 mM EDTA), and twice with TE (10 mM Tris-Cl pH 8, 1 mM EDTA). DNA was eluted with 100 µl elution buffer (1% SDS, 10 mM EDTA pH 8, 50 mM Tris-Cl pH 8) for 15 min at 65 °C after vigorous shaking. Supernatant was recovered with centrifugation (1 min, RT, 11000x*g*), beads washed with 150 µl elution buffer and pooled together. DNA was decrosslinked overnight at 65 °C in in presence of 1 µg Proteinase K and isolated with Phenol:Chloroform (Sigma, Cat#P2069) extraction and ethanol precipitation. Sequencing libraries were prepared using NEBNext Ultra II DNA Library Prep kit (NEB, Cat#E7645L) following the manufacturer's protocol and in-house adaptors with 8 bp UMIs in addition to unique barcodes. Libraries were sequenced on NextSeq 500 or 2000 either with single or paired-end mode for at least 50 cycles.

**ChIP-qPCR:** ES cell chromatin was prepared and quantified as above. Equal amounts of DNA were used for ChIP, DNA was isolated and qPCR was performed for both input (1:20 dilution) and ChIP samples. qPCR was performed in 15 µl volume with LightCycler 480 SYBR Green I Master (Roche, Cat#4887352001). Oligos used were as follows:

GTATACCTCTAACTGCCACAAGT and TCAATGGGGAACTAATGGAAAGA,

ACAGGGTTTGAGCTACAGTTAA and TGGGATTCTCTACTTTGCAGATG.

**Spike-in CUT&Tag:** To perform R-loop CUT&Tag in ES cells in the presence of MCF-7 cells, 100,000 ES cells were first bound to 10 µl concanavalin A beads (Epicypher, Cat#21-1411) per condition with binding buffer (20 mM HEPES pH 7.5, 10 mM KCl, 1 mM $CaCl_2$, 1 mM $MnCl_2$). To deplete R-loops, ES cells bound to 10 µl beads were dissolved in 100 µl RNase H reaction buffer and incubated with 10 µl RNase H (NEB, Cat#M0297S) at 37 °C for 2 h. As a control, RNase H enzyme was skipped. After 2 h incubation, beads were washed thrice with wash buffer (20 mM HEPES pH 7.5, 150 mM NaCl, 0.5 mM spermidine, protease inhibitors) to remove RNase H and dissolved in 100 µl antibody buffer (wash buffer with 2 mM EDTA pH

8, 0.01% BSA, 0.05% digitonin, protease inhibitors). In parallel, 100,000 MCF-7 cells were bound to 10 µl beads and dissolved in 100 µl antibody buffer. 10 µl if this mix (10,000 MCF-7 cells) were mixed with 100,000 ES cells bound to beads in 100 µl antibody buffer for each condition. Beads were incubated overnight at 4 °C with 1 µl S9.6 antibody (Sigma, Cat#MABE1095). Next day, beads were recovered and incubated with 1 µL secondary antibody (Epicypher Cat#13-0048) in 100 µL in Dig-wash buffer (20 mM HEPES pH 7.5, 150 mM NaCl, 0.5 mM spermidine, 0.05% digitonin, protease inhibitors) for 1 h at RT. Beads were washed thrice with 1 mL Dig-wash buffer and incubated with 50 µl of pA-Tn5 mix (2.5 µL pA-Tn5 adapter complex (Epicypher, Cat#15-1017) in 50 µL Dig-300 buffer) for 1 h at RT. Beads were washed thrice with Dig-300 buffer (20 mM HEPES pH 7.5, 300 mM NaCl, 0.5 mM spermidine, 0.01% digitonin, protease inhibitors) and Tn5 was activated with 300 µL Tagmentation buffer (Dig-300 buffer with 10 mM $MgCl_2$) for 1 h at 37 ºC. To stop tagmentation, beads were incubated with 10 µL 0.5 M EDTA, 1.5 µL 20% SDS and 2.5 µl Proteinase K (20mg/ml) for 30 min at 37 ºC and then 50 ºC for 1 h. DNA was purified by adding 300 µl phenol chloroform mix, transferred to phase lock tube (Qiagen, Cat#129046) and centrifuged (5 min, RT, 16000x*g*). 300 µl Chloroform: Isoamyl alcohol mix (Sigma, Cat#25666) was added to tubes and centrifuged again (5 min, RT, 16000x*g*). Aqueous phase was precipitated with 100% ethanol, washed with 80% ethanol and dissolved in 25 µl TE buffer with 1/400 RNase A (Thermo Fisher, Cat#EN0531). DNA was incubated for 1 h at 37 °C. PCR was performed with 21 µl tagmented DNA in presence of 0.4 µM universal i5 primer, 0.4 µM barcoded i7 primers and 25 µL NEBNext HiFi 2x PCR Master mix (NEB, Cat#M0541S). PCR cycles were as follows: 72 °C for 5 min, then 10 cycles of 98 °C for 40 sec, 63 °C for 10 sec and final cycle of 72 °C for 1 min. DNA was purified with 1.1x SPRI beads and eluted in 25 µL 0.1X TE buffer. Libraries were sequenced as above.

**Spike-in poly-A RNA-seq:** To measure gene expression following GapR induction, ES cells harboring GapR were induced overnight with 1 µg/ml dox. Uninduced and induced ($10^6$) GapR cells were harvested in parallel with standard trypsin dissociation, collected by centrifugation in a 1.5 ml eppendorf containing 0.5 x $10^6$ *Drosophila* S2 cells. Pellet was dissolved in 1 ml TRIzol (Thermo Fisher, Cat#15596026) and flash frozen in liquid $N_2$. Poly (A)

RNA isolation, RNA quality control, strand-specific library preparation and 100 bp paired-end sequencing was performed at CeGaT GmbH, Germany, using TS Flex mRNA protocol for at least 100 million reads.

**Bioinformatic analyses:** BCL files were demultiplexed and converted to fastq format with no barcode mismatches using bcl2fastq v2.20.0. Reads were aligned independently to mm10 genome for mouse, hg38 for human, dm6 for *Drosophila* and MG1655 (build 2001-10-15) for *E. coli* using Bowtie2 v2.3.5.1 (Langmead and Salzberg 2012) in local mode with --very-sensitive-local, -X 1000, -I 0 options and bam files were created using samtools v1.16. Duplicates reads were removed from bam files using UMIs for ChIP-seq reads and with Picard v2.23.3 for CUT&Tag reads, in addition to filtering reads with MPAQ <30. Reads mapping to blacklist regions were removed and bigwig files were generated with deepTools (Ramírez et al. 2016) bamCoverage v3.4.1 by multiplying reads in mouse genome to 1/number of reads mapping to either *E. coli*, human or *Drosophila* genomes for each replicate. RNA-seq reads were processed with nf-core/rna-seq v3.14.0 using gtf gencode M25. Reads trimmed with Trim Galore, aligned with STAR v2.7.10a and count table was generated using SALMON v1.10.1.

Peaks were called on each replicate with MACS v3.0.1 callpeak option considering input files and FDR 0.05 (Zhang et al. 2008). Next, peaks were called on merged replicates with FDR 0.01. Peaks arising from merged replicates needed to be present in at least 2 out of 3 replicates to be considered in the final peak set. For GapR, we called broad peaks with --max-gap 300 and FDR 0.01 options. Peaks were annotated with ChIPseeker v1.42.1 (Yu et al. 2015) using Bioconductor package TxDb.Mmusculus.UCSC.mm10.knownGene v.3.10.0 and with ChromHMM states (Pintacuda et al. 2017) for mm10 genome with OverlapEnrichment command.

Metaplots were generated using computeMatrix scale-regions or reference-point options, followed by plotProfile or plotHeatmap commands. Final tables were imported and plotted in R v4.4.3. Venn diagrams were created with VennDiagram package v1.7.3. Genome browser snapshots were generated using python package SparK with option -sm 10

(Kurtenbach and William Harbour 2019). To quantify signal at a given region, bigWigAverageOverBed UCSC-tools was used for defined windows for each replicate separately. Signal was averaged over replicates and z-score normalized in R. Figures were prepared using ggplot2 library. Central line in boxplots represent the median, box represents the interquartile range and whiskers extend to the values within 1.5x IQR from Q1 and Q3.

Following tables and datasets were downloaded and processed, if required, as described above: Active and Primed enhancers (Cruz-Molina et al. 2017), Super enhancers (Novo et al. 2018), Micro-C loops (Hsieh et al. 2022), BisMapR R-loops (GSE160578; (Wulfridge and Sarma 2021)), Rad21 ChIP-seq (GSE135180; (Liu et al. 2021)) and Smc1 ChIP-seq (GSE131356; (Owens et al.)), post mitotic release RNA-seq (GSE196889; (Chervova et al. 2023)).
